# Structural insights into distinct signaling profiles of the μOR activated by diverse agonists

**DOI:** 10.1101/2021.12.07.471645

**Authors:** Qianhui Qu, Weijiao Huang, Deniz Aydin, Joseph M. Paggi, Alpay B. Seven, Haoqing Wang, Soumen Chakraborty, Tao Che, Jeffrey F. DiBerto, Michael J. Robertson, Asuka Inoue, Bryan L. Roth, Susruta Majumdar, Ron O. Dror, Brian K. Kobilka, Georgios Skiniotis

## Abstract

Drugs targeting the G protein-coupled μ-opioid receptor (μOR) are the most effective analgesics available but are also associated with fatal respiratory depression. While some partial opioid agonists appear to be safer than full agonists, the signaling pathways responsible for respiratory depression have yet to be elucidated. Here we investigated the structural and mechanistic basis of action of lofentanil (LFT) and mitragynine pseudoindoxyl (MP), two μOR agonists with different safety profiles. LFT, one of the most potent and lethal opioids, and MP, a derivative from the kratom plant with reduced respiratory depression in animal studies at equianalgesic doses, exhibited markedly different signaling efficacy profiles for G protein subtype activation and recruitment of β-arrestins. Cryo-EM structures of the μOR-Gi1 complex with MP (2.5Å) and LFT (3.2Å) revealed that the two ligands engage distinct sub-pockets, and molecular dynamics (MD) simulations showed additional differences in the binding site that propagate to the intracellular side of the receptor where G proteins and β-arrestins bind. While MP favors the precise G protein-bound active state observed in the cryo-EM structures, LFT favors a distinct active state. These results highlight how drugs engaging different parts of the μOR orthosteric pocket can lead to distinct signaling outcomes.

## Introduction

Opioids targeting the μ-opioid receptor (μOR), such as the natural alkaloid morphine and synthetic agonists like fentanyl, remain the most effective analgesics for treating acute and chronic pain. The μOR is a G protein-coupled receptor (GPCR) that signals through six different heterotrimeric G protein subtypes: Gi1, Gi2, Gi3, GoA, GoB, and Gz (henceforth the Gi/o/z family). However, μOR activation can also recruit β-arrestins, which not only promote receptor endocytosis but also drive G protein-independent signaling.

Opioid receptor ligands range from small synthetic molecules to plant alkaloids and peptides with diverse scaffolds and distinct signaling properties^1^. Fentanyl and a series of congeners have been synthesized to initiate strong and rapid analgesia, and are commonly prescribed to treat chronic cancer pain or used in anesthesia management^2^. Although the pharmacology of these broadly used fentanyl compounds is well characterized, their detailed interaction network with the μOR has not been determined. Fentanyl analogues demonstrate an enhanced ability to desensitize μOR^3^, and preferentially recruit β-arrestin-2 in PathHunter cellular assays^4^. LFT, in particular (Fig. 1a), is one of the most potent fentanyl analogues. At over 10,000 times more potent than morphine^5^, LFT has an increased risk of addiction and overdose, and is therefore not used clinically.

**Fig. 1.**
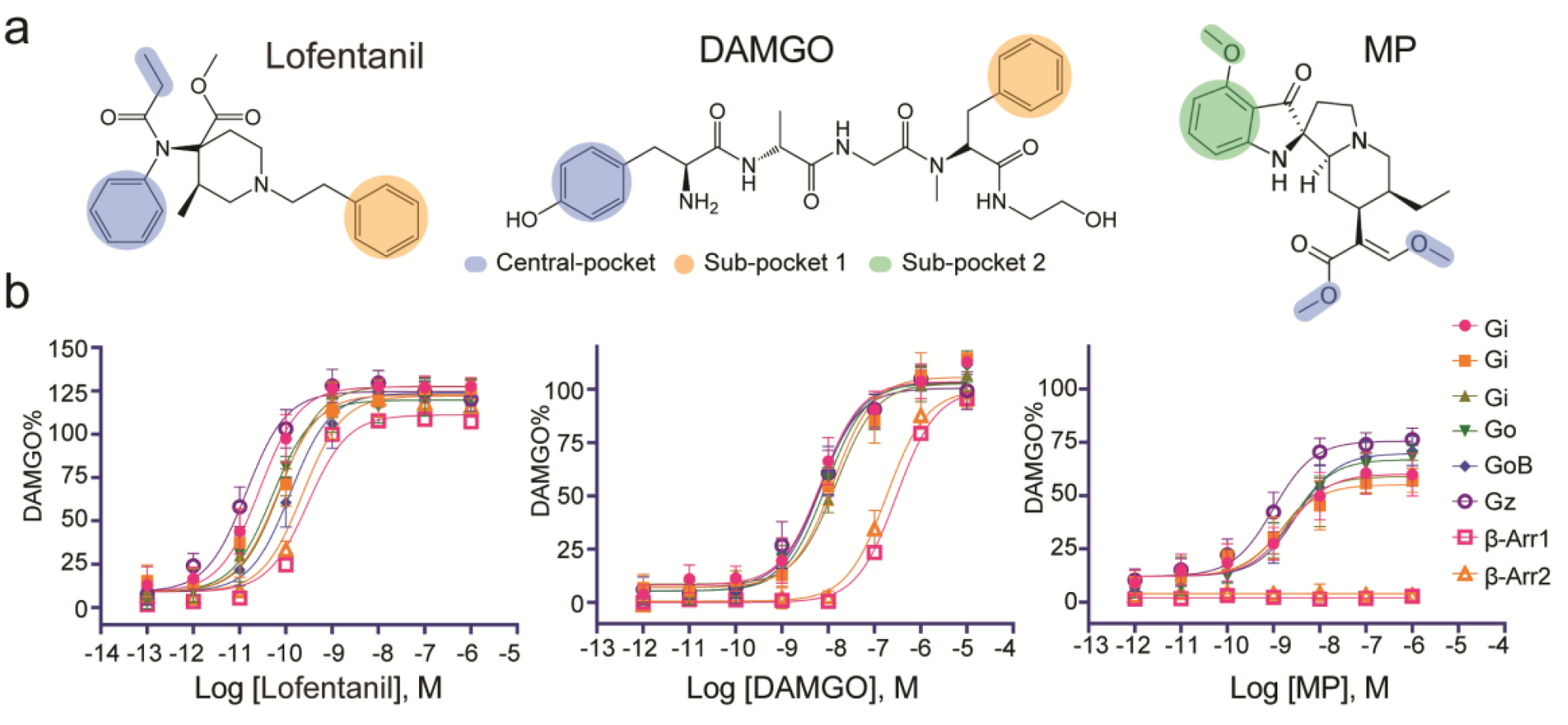
Biased and non-biased ligands for μOR. **a**, Structurally diverse μOR ligands with distinct pharmacological properties. DAMGO, (D-Ala^2^, N-MePhe^4^, Gly-ol)-enkephalin; Lofentanil; MP, Mitragynine pseudoindoxyl. **b**, Concentration-dependent activation of Gi1, Gi2, Gi3, GoA, Go and Gz, and recruitment of β-arrestin-1 and β-arrestin-2 using BRET-based biosensors. Data for all functional assays that were carried out in hMOR were normalized to *E*_max_ of DAMGO. The dose response curves were fit using a three-parameter logistic equation in GraphPad Prism. See also Extended Data Fig. 1 and Extended Data Table 1 for efficacy and potency data for ligands.

On the other hand, kratom-derived MP (Fig. 1a) is an indole-based analgesic alkaloid that is structurally different from the endogenous enkephalin-derived peptides, morphine derivatives, and synthetic analogs based on fentanyl (Fig. 1a). Kratom extract from the leaves of the tropical evergreen tree *Mitragyna speciosa* found in Southeast Asia has been used for centuries by local cultures to enhance endurance and combat fatigue. Recently, kratom has gained global popularity for its ability to relieve pain and alleviate symptoms during opioid withdrawal, and it is also used recreationally with a claimed reduced addiction liability^6,7^. There are nearly 54 alkaloids present in kratom with mitragynine being the major alkaloid (∼66% of total alkaloid content). It is currently believed that the μOR dependent oral analgesic actions of kratom are derived from the metabolism of mitragynine to 7-hydroxy mitragynine (7OH)^8^. Recent reports have also shown that MP is a minor metabolite of mitrgynine^9,10^. Previously, we synthesized and systematically examined the pharmacological behaviors of a series of mitragynine-based natural products and analogs that included mitragynine, 7OH, and MP both *in vitro* and *in vivo*. We found that, mice treated with MP developed antinociceptive tolerance much more slowly compared with morphine, and that MP did not cause respiratory depression at doses that produce analgesia comparable to morphine at μOR^11^.

The unwanted effect of respiratory depression was previously attributed to μOR signaling through β-arrestins^12,13^, fueling efforts to discover biased agonists that selectively stimulate G protein over arrestin^14-16^. However, a number of studies have raised doubts about the role of arrestin signaling in respiratory depression^17-20^, and recent work argues for balanced agonists as a mechanism to circumvent tolerance mediated by opioids^21^. On the other hand, there are conflicting reports on the role of specific Gi/o/z subtypes in the actions of opioids *in vivo*. For instance, experiments using antisense RNA to reduce the expression of Gαi1, Gαi2 or Gαi3 showed an impaired supraspinal analgesic response to morphine only in mice treated with antisense RNA to Gαi2^22^. Of interest, analgesia produced by sufentanil in this study remained intact in mice treated with antisense RNA to Gαi2. However, these results conflict with other studies showing that supraspinal analgesia was intact in Gαi2 and Gαi3 knockout mice, whereas an impaired analgesic response to morphine was observed in heterozygous Gαo knockout mice^1^. In contrast, Gαz KO mice showed little or no change in the supraspinal analgesia of a single dose of morphine, yet there was a marked increase in analgesic tolerance, and a decrease in lethality where the LD_50_ (dose at which 50% of animals die) was 700 mg/kg for wildtype and greater than 800 mg/kg for homozygous Gαz knockout mice^23^. Notably, none of the studies above specifically examined respiratory depression, which is a frequent cause of morbidity and mortality among opioid users. It is thus currently unclear, whether analgesia and/or respiratory depression may be mediated by more than one Gi/o/z subtype in an agonist specific manner.

To explore the molecular mechanisms contributing to μOR mediated respiratory depression, we examined the transducer coupling propensities along with the structural effects of LFT and MP which differ in their *in vivo* potency and tendency to promote respiratory depression at equianalgesic doses. We observed that LFT and MP have distinct efficacy signaling profiles at Gi1, Gi2, Gi3, GoA, GoB and Gz, as well as their recruitment of β-arrestins, with structural analysis of the μOR-Gi1 complex bound to each of these ligands providing a glimpse into unique allosteric pathways that may contribute to these drugs’ differential signaling profiles.

## Results

### Distinct signaling profiles for μOR activated by LFT and MP

Given the diametrically opposite attributes of LFT and MP for safety and potency, we performed a detailed comparison of their signaling profiles using the TRUPATH bioluminescence-resonance energy (BRET) platform^24^ for Gi1, Gi2, Gi3, GoA, GoB and Gz activation, and a complimentary BRET-based β-arrestin-1 and -2 recruitment assay (Fig. 1b). The efficacies for MP and LFT were expressed as a percentage of the response to the reference ligand DAMGO, a peptide analog of the endogenous opioid met-enkephalin. DAMGO which had a narrow, ∼2-fold potency spread with highest activity at Gz and lowest at Gi3, showed an observed potency rank order of Gz∼Gi1>GoB∼GoA>Gi2∼Gi3 (Extended Data Table 1).

Compared to DAMGO, LFT promotes activation of all Gi/o/z proteins with higher efficacy at all subtypes, while arrestins are recruited to a comparable or greater level. The potency of activation of Gz was the highest, at more than nine times greater than activation of Gi3 and a potency rank order of Gz>Gi1>GoA∼Gi2∼Gi3>GoB. (Fig. 1b, Extended Data Table 1). In contrast, MP was a partial agonist at all Gi/o/z subtypes, with no detectable recruitment of β-arrestins. The potency of MP was similar for all six G protein subtypes with a potency spread of ∼3-fold between Gz (0.9 nM) and GoB (3.2 nM); however, the efficacy at all Gi/o/z and arrestin subtypes was lower than DAMGO. Notably, the efficacy of MP against DAMGO and LFT as well as morphine at the three most abundant Gα-subtypes present in the brain (i.e., Gz, GoA, GoB and Gi1) was found to be significantly lower (Fig. 1b, Extended Data Fig. 1). Another striking difference between MP and LFT is recruitment of β-arrestin-1 and β-arrestin-2, which is almost undetectable for MP in the BRET recruitment assay (Fig. 1b). The relative potencies and efficacies for Gi1, β-arrestin-1 and β-arrestin-2 were confirmed using Nano-BiT^25^ enzyme complementation assays that monitored G protein dissociation and arrestin recruitment (Extended Data Fig.1). As expected, LFT preferentially promoted the most efficacious β-arrestin-1 and -2 recruitment, while it was slightly more efficacious at Gi heterotrimer protein dissociation. In contrast, MP exhibited very little detectable activity towards recruitment of β-arrestins, and was as potent as DAMGO in G protein dissociation assays, consistent with the TRUPATH data (Fig. 1b) and previous characterization by GTPγS binding^11^ and cAMP^26^ assays.

### Structures of μOR-Gi1 complex with MP or LFT

To probe the molecular basis of different signaling behaviors of μOR modulated by MP and LFT, we obtained cryo-EM structures of MP-μOR-Gi1 and LFT-μOR-Gi1-scFv complexes, as Gi1 is in the middle of the potency profiles for both MP and LFT (Fig. 1b). The map for LFT-μOR-Gi1-scFv was determined from holey carbon grids at a global nominal resolution of 3.2Å, whereas the map for MP-μOR-Gi1 was initially globally determined to 2.5Å, owing primarily to the use of holey gold grids that minimized specimen motion during data collection. Notably, local refinement followed by density modification in Phenix, further improved the resolution of the map in several regions, revealing a number of well resolved water molecules (Figs. 2a-b and Extended Data Figs. 2-3). The high-quality density maps enabled *de novo* modeling of ligands MP and LFT, and the atomic coordinates were further optimized using GemSpot^27^ (Figs. 2a and 2b insets). The overall architectures of both MP-μOR-Gi1 and LFT-μOR-Gi1 are similar to the enkephalin-like agonist DAMGO-bound μOR-Gi1 structure^28^ (Extended Data Fig. 4a). General structural hallmarks of GPCR activation, including the DR^3.50^Y, CW^6.48^xP^6.50^, P^5.50^-I^3.40^-F^6.44^ and NP^7.50^xxY^7.53^ motifs (superscripts denote generic Ballesteros–Weinstein numbering^29^) are essentially identical to that of DAMGO^30^ and morphinan agonist BU72^31^, with the latter being co-crystallized with a G protein mimetic nanobody (Extended Data Fig. 4b-d). Also very similar is the interface between nucleotide-free G protein and μOR in the MP, LFT and DAMGO structures, representing a canonical μOR-Gi1 coupling state (Extended Data Fig. 4e,f).

**Fig. 2.**
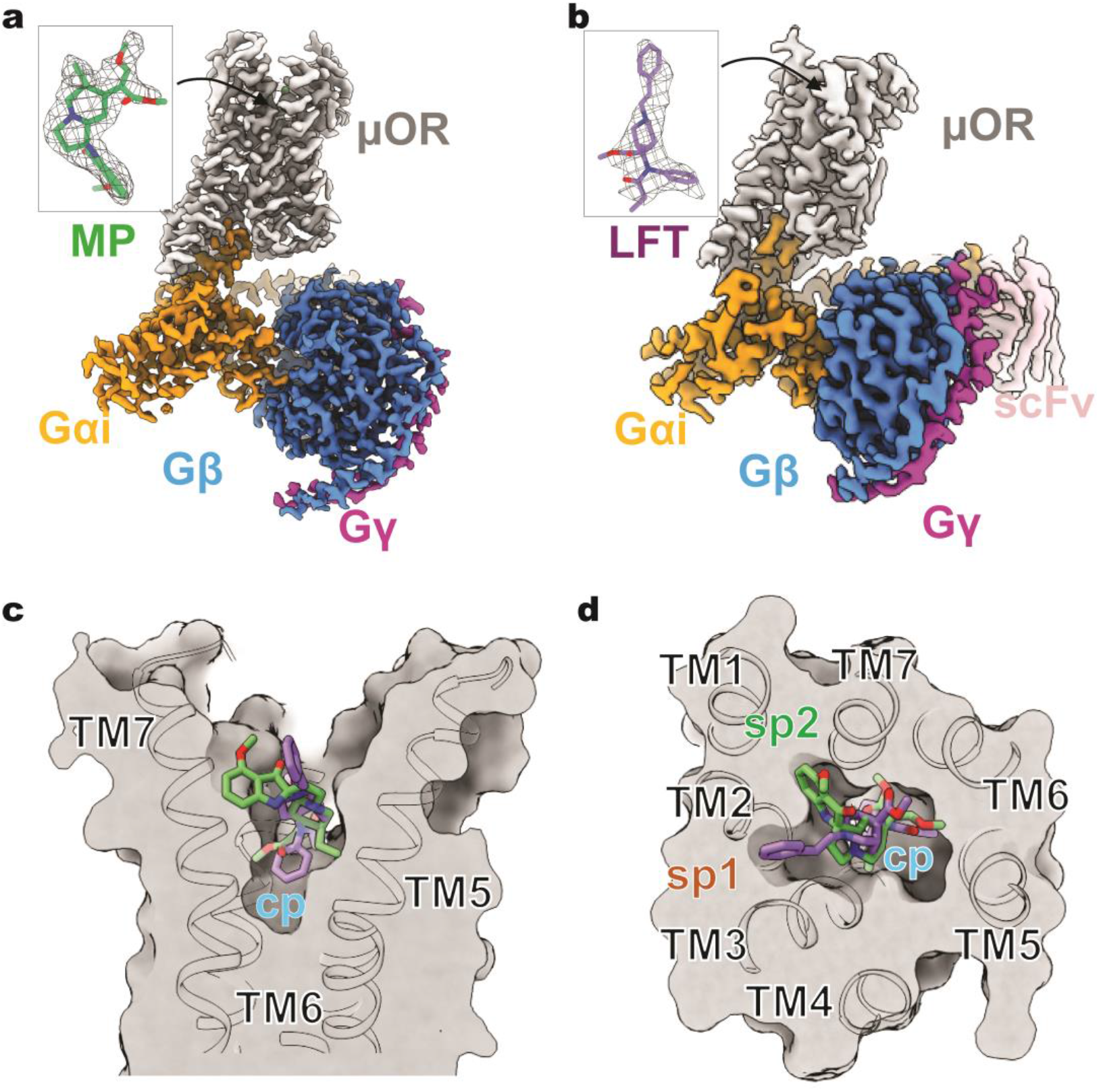
Structures of μOR-Gi complex activated by Gi1-biased partial agonist MP and arrestin-biased full agonist LFT. **a**,**b**, 2.5Åcryo-EM map for the MP-μOR-Gi1 complex (**a**) and 3.2Åcryo-EM map for the LFT-μOR-Gi1-scFv complex (**b**). The insets highlight the well-resolved density (shown as wire-net) for MP (green) and LFT (purple). μOR is colored grey, Gαi1 in orange, Gβ in dodger blue, Gγ in magenta and scFv in pink. **c**,**d**, Superposition between the MP-μOR-Gi and LFT-μOR-Gi structures shows that the structurally distinct MP and LFT occupy both a common central pocket (cp) in the orthosteric binding site (**c**, viewed from membrane plane, **d**, viewed from extracellular side), while occupying different sub-pockets (sp1 and sp2).

Notwithstanding the diverse structural scaffolds, both MP and LFT fit snugly in the same orthosteric site in the μOR composed of the extracellular side on transmembrane (TM) helices 2,3, 5, 6 and 7, which is also occupied by DAMGO, BU72 and the covalent antagonist βFNA^32^ (Fig. 2c, d and Extended Data Fig.5). The elongated LFT has an orientation similar to DAMGO and BU72, with the hydrophobic 1-phenethyl branch of LFT pointing towards a sub-pocket (sp1) formed by TM2-3 and ECL1-2, which is occupied by the bulky phenyl group of BU72 and the methyl-phenylalanine of DAMGO (Extended Data Fig.5). In contrast, in addition to the central pocket (cp) shared by all ligands, MP occupies a novel sub-pocket (sp2) formed by TM1, TM2 and TM7, (Fig. 2d and Extended Data Fig.5b). Of interest, the 9-methoxy group on the indole ring orients towards the spacious μOR extracellular outlet, which may explain the observation that various substituents on this indole C-9 position could retain similar affinity for the μOR^11^. We also note that judging from the suboptimal fit of the β-methoxyacrylate tail of MP in the cryoEM density, this moiety may interchangeably adopt two diametrically opposite orientations (Extended Data Fig. 3b).

The fentanyl derived synthetic compounds share a piperidinyl core and exhibit various *in vivo* potencies for the μOR, with fentanyl being approximately 80-100 times more potent than morphine in rodents. Addition of a carbomethoxy moiety onto the 4-axial position of the fentanyl piperidinyl group makes carfentanil nearly 10,000 times more potent than morphine (Extended Data Fig.6a). To further probe the binding properties of fentanyl synthetics, we employed the Maestro docking software (Schrödinger Release 2018-4: Maestro, Schrödinger, LLC, New York, NY, 2018) to generate fentanyl and carfentanil models based on the LFT coordinates within the LFT-μOR cryo-EM structure. As expected, carfentanil adopts a similar configuration as LFT. On the other hand, docking generated two high-scoring poses for fentanyl assuming starkly opposite orientations (Extended Data Fig.6b-d). To address this discrepancy, we employed molecular dynamics (MD) simulations to assess the stability of each pose.

The simulations showed that a pose analogous to the cryo-EM pose of LFT is more stable than either of the fentanyl poses obtained through docking. (Extended Data Fig.6e), thus suggesting that fentanyl tends to adopt an LFT-like pose.

### Ligand binding pocket interactions and dynamics

Closer inspection of the ligand binding network highlights a highly conserved polar interaction between μOR D147^3.32^ and a protonatable amine (NH^+^) of μOR ligands (Fig. 3 and Extended Data Fig.5c). Previous μOR structures have shown that the phenol groups of both DAMGO and BU72 are positioned in the central pocket with their hydroxyl groups oriented to form a water mediated interaction with H297^6.52^. Here we observe that LFT and MP position their aniline and carbomethoxy groups, respectively, deeper into the central pocket (Fig.3, residues shown as grey sticks), forming extensive contacts with M151^3.36^, W293^6.48^, I296^6.51^, I322^7.39^ and Y326^7.43^. In lieu of the phenolic hydroxyl group of DAMGO and morphinan ligands, LFT and MP position their N-propylamide and methoxyenolate moieties, respectively, in the TM5-6 region.

**Fig. 3.**
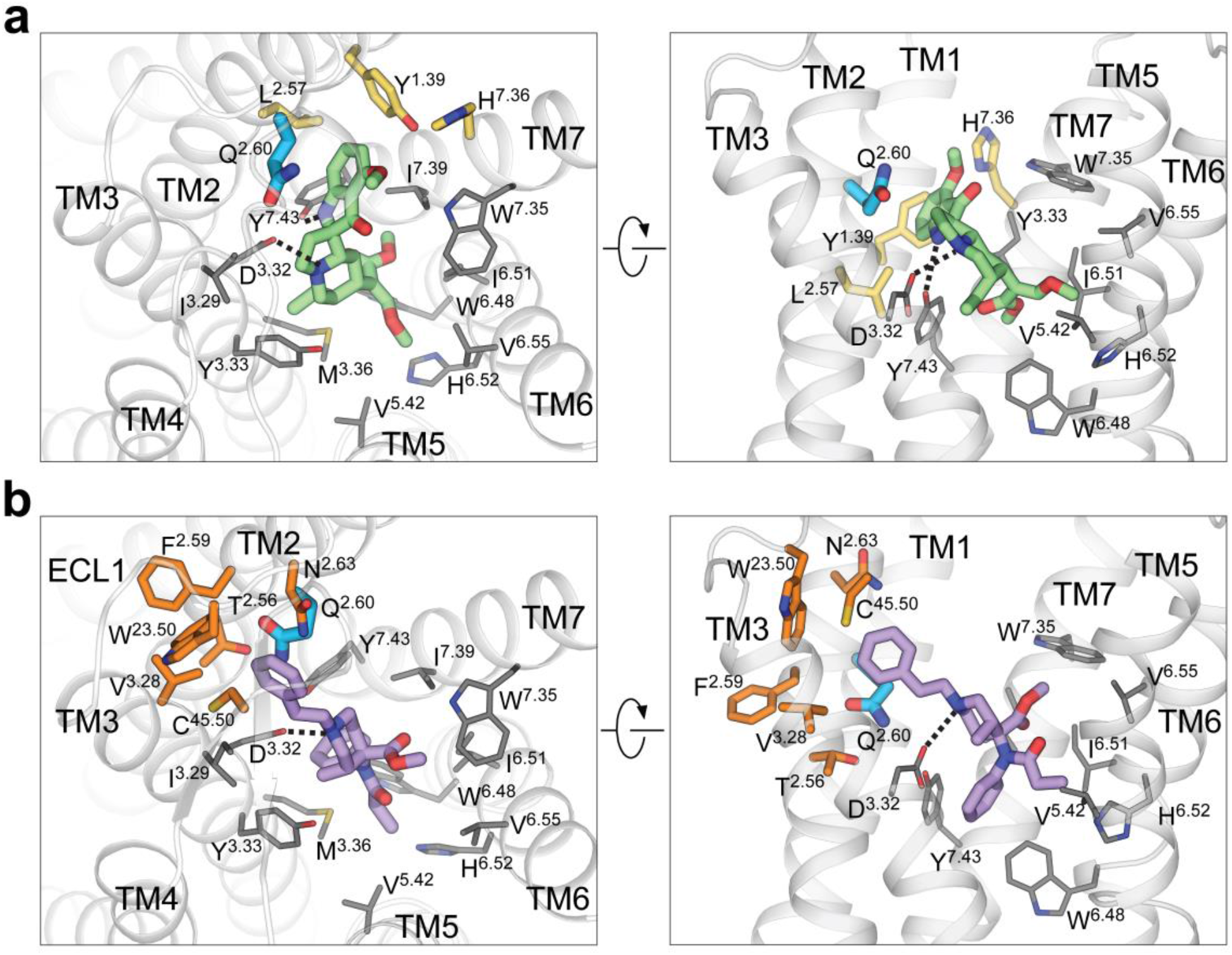
Semi-conserved ligand interaction network for MP and LFT. **a**,**b**, The μOR orthosteric binding pocket for MP (**a**) and LFT (**b**) viewed from the extracellular side (left panels) and membrane plane (right panels). Residues involved in both MP and LFT interaction are colored grey, while residues uniquely contributing to MP interaction are shown in yellow, and those for LFT in orange. Residue Q124^2.60^, which interacts with both MP and LFT but in different orientations, is highlighted in blue. Dashed lines depict polar interactions.

The side-chain of residue Q124^2.60^ (blue stick, Fig.3) separates sub-pockets sp1 and sp2 (represented by orange and yellow sticks, respectively, in Fig. 3) with a different side-chain orientation in the MP-bound structure, allowing space for the indole ring of MP. The interaction between the indole ring and Q124^2.60^ is shown to be critical for MP agonism, as the μOR Q124^2.60^A mutant reduces by more than half the maximal Gi response to MP and completely abolishes arrestin recruitment (Extended Data Fig. 7). This interaction is less dominant in other ligands; the Q124^2.60^A mutant retains full Gi response to LFT and more than 70% of the maximal Gi1 response to DAMGO, although potency is reduced by more than 100-fold for DAMGO (Extended Data Fig.7). Using MD simulations, we found that when not sterically blocked, Q124^2.60^ can form a hydrogen bond to Y326^7.43^ stabilizing it in an inward position. In simulations with LFT, this interaction is nearly always present (Fig.4a,b). Interestingly, we found that DAMGO induces this interaction only part of the time. The difference in stability of this interaction for LFT and DAMGO can largely be explained by a difference in positioning of Y128^2.64^. In simulations with LFT, Y128^2.64^ forms a tight interaction with Q124^2.60^, thereby stabilizing an interaction with Y326^7.43^ (Fig.4a,b). However, this interaction rarely forms in simulations with DAMGO, likely due to interference from the peptide backbone of the ligand.

**Fig. 4.**
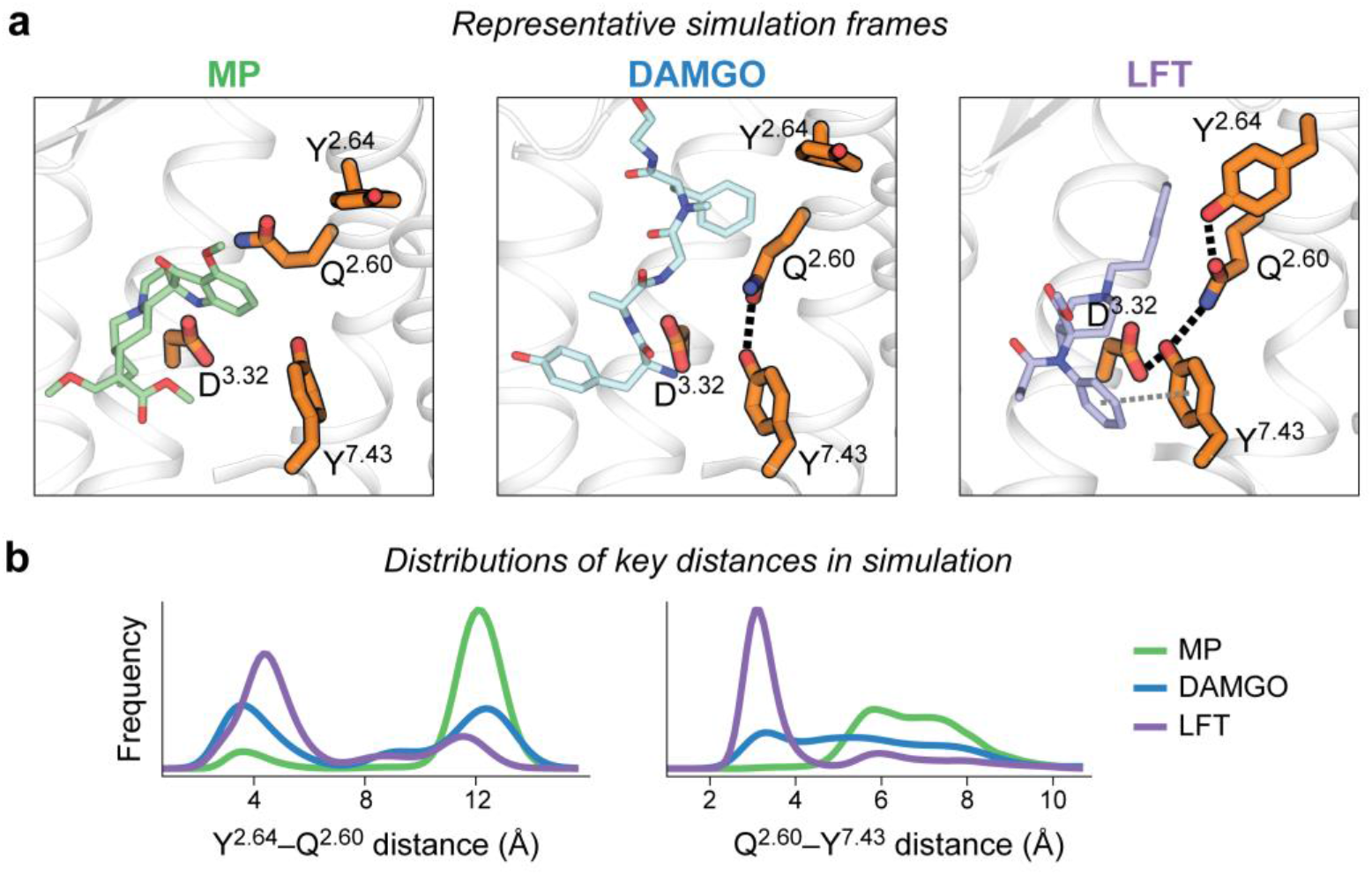
Simulations of the μOR reveal distinct binding pocket conformations favored by MP, DAMGO, and LFT. **(a)** Representative simulation frames showing ligand-specific hydrogen bonding networks in the binding pocket, involving key amino acid residues Y326^7.43^, Q124^2.60^, Y128^2.64^, and D147^3.32^. **(b)** Distributions of key inter-residue distances in simulations with various ligands bound (see Methods). MP and LFT favor extreme conformations, whereas DAMGO samples a broader range of conformations, including the two extreme conformations favored by MP and LFT. Black dashed lines represent hydrogen bonding interactions between the key residues, and the grey dashed line represents the π–π interaction between Y326^7.43^ and the aniline group phenyl ring of LFT. Simulations were performed without a bound G protein.

The MP-bound structure shows a hydrogen-bond formed between Y326^7.43^ and the indole amine (N1) of MP (Fig. 3a). The N1 is critical for MP activity as alkyl substitution reduces its receptor affinity^11^. In simulations, stable π-π stacking is observed between LFT’s aromatic aniline ring and Y326^7.43^. Together with the strong Y326^7.43^–Q124^2.60^ interaction, this allows for more inward conformations of Y326^7.43^ in LFT bound μOR (Fig. 4, Extended Data Fig. 8). In contrast, the hydrogen-bonding between Y326^7.43^ and MP is less stable, as the direct hydrogen bond can be replaced by a water-mediated interaction. Even though the hydrogen bond between Y326^7.43^ and DAMGO is also liable to be replaced by a water-mediated interaction, the Y326^7.43^–Q124^2.60^interaction, which is often formed in the presence of DAMGO, results in an intermediate positioning of Y326^7.43^ (Fig. 4, Extended Data Fig. 8). These differences in the dynamics of Y326^7.43^ result in LFT favoring more counterclockwise rotations of TM7 (viewed from the extracellular side) when compared with MP. As described below, our simulations suggest that this rotation of TM7 near the ligand binding pocket influences the conformation of the receptor’s intracellular surface.

### Allosteric effects of ligands on the transducer interface

It has been hypothesized that different ligands modulate the equilibrium among multiple conformations of μOR, which in turn favor signaling through different transducers^33^. For μOR, it was not previously known what these conformations are, nor how different ligands select between them. The structures reported here, along with our previous structure of μOR bound to DAMGO, illuminate differences in protein– ligand interactions for ligands with a range of distinct pharmacological profiles. However, by themselves, these structures do not reveal how different binding pocket conformations result in distinct intracellular conformations, likely because the presence of a G protein overwhelms the effect of the ligands on the conformation of the receptor.

To understand how differences in the ligands propagate to the intracellular transducer binding site, we performed atomistic simulations with the G protein removed. We initiated simulations of μOR bound to MP, LFT, and DAMGO from the structures presented in this manuscript and the previously published DAMGO-bound μOR structure, respectively. We hypothesized that in the absence of a G protein, the receptor would relax away from the G protein–bound state observed in the cryo-EM structures, revealing ligand-dependent differences in the conformational ensemble of the receptor.

The simulations showed two major conformational states of the intracellular coupling site that differ in TM7 (Fig. 5). One state, which we refer to as the “canonical active state”, is essentially identical to the intracellular conformation observed in the G protein–bound cryo-EM structures. The other state, which we refer to as the “alternative state”, differs in that TM7 is rotated counterclockwise (viewed from the extracellular side), with the intracellular portion of TM7 positioned inwards (towards TM3) and the kink in the NPxxY region relaxed (Fig. 5a,b). Notably, MP favors the canonical active state, LFT favors the alternative state, and DAMGO favors an equilibrium between these two states (Fig. 5d). These conformations may provide a link to the distinct transducer recruitment profiles reflected in our TRUPATH assays.

**Fig. 5.**
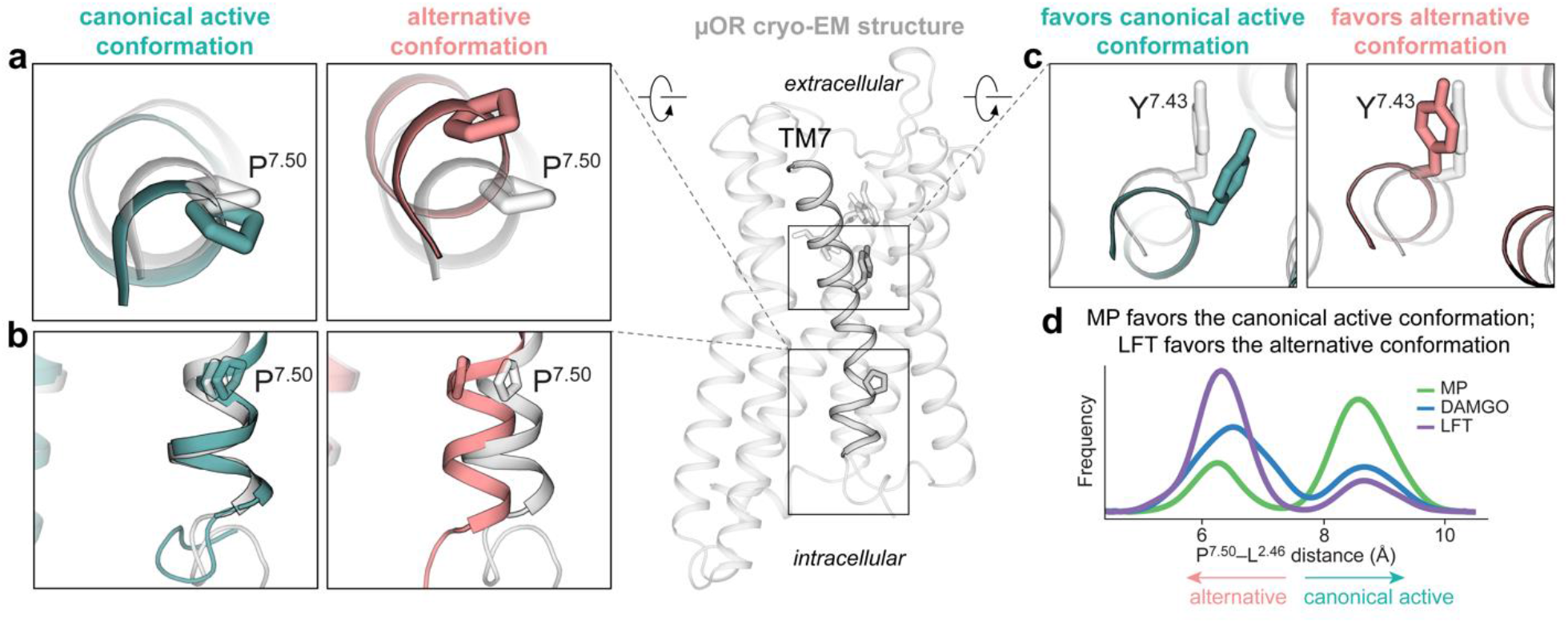
In simulations with the G protein removed, the μOR adopts two active intracellular conformations, with MP and LFT favoring different conformations. **(a, b)** At the intracellular coupling site, the canonical active conformation matches the intracellular conformation observed in the G protein–bound cryo-EM structures. The counterclockwise twist at TM7 during the transition from the canonical active conformation to the alternative conformation triggers an inward movement of P333^7.50^. The grey rendering is based on the MP-μOR-Gi cryo-EM structure, but the TM7 structural features shown are essentially identical in the MP-μOR-Gi, LFT-μOR-Gi, and DAMGO-μOR-Gi cryo-EM structures. **(c)** A clockwise rotation of TM7 at the ligand binding site, specifically at Y326^7.43^, favors the canonical active conformation, whereas a counterclockwise rotation favors the alternative state. **(d)** Relative to DAMGO, MP favors the canonical active conformation, whereas LFT favors the alternative conformation.

The ligands control the occupancy of these two states by favoring different rotations of TM7 (Fig. 5c). LFT is associated with a counterclockwise rotation of TM7, whereas MP is associated with a clockwise rotation of TM7. This rotation of TM7 is enabled by a change in the hydrogen bonding network in the sodium binding pocket. In the canonical active state, hydrogen bonds are present between N86^1.50^–S329^7.46^ and D114^2.50^– N332^7.49^. In the alternative state, these interactions are broken and replaced by hydrogen bonds between D114^2.50^–S329^7.46^ and D114^2.50^–N150^3.35^ (Extended Data Fig. 9). MP and LFT favor intracellular μOR conformations that are similar to those observed at the angiotensin II type 1 receptor (AT_1_R) for G-protein-biased and arrestin-biased ligands, respectively^34^ (Extended Data Fig. 10a). For both μOR and AT_1_R, the transition from the canonical active conformation to the alternative conformation involves a counterclockwise twist at TM7, leading to the inward movement of P333^7.50^ and the shift of the hydrogen bonding network in the sodium binding pocket. The inward TM7 in the alternative conformation is farther from the inactive state for both receptors (Extended Data Fig. 10a). We also note that the alternative conformations of μOR and AT_1_R differ in that the intracellular end of TM7 is inwardly displaced in μOR, discouraging the downward Y336^7.53^ rotamer observed at AT_1_R (Extended Data Fig. 10b). This appears to be due to the interaction between R165^3.50^ and two negatively charged residues at helix 8 (D340^8.47^ and E341^8.48^) that are unique to opioid receptors.

To validate the role of this ligand-stabilized rotation of TM7 in determining signaling behaviors, we characterized the signaling profile of MP, DAMGO, and LFT for the Y326^7.43^F mutant. Since the loss of the hydroxyl group would disrupt the hydrogen bonds to Q124^2.60^ and D147^3.32^ (Fig. 4a), we hypothesized that this mutation would favor outward conformations of Y326^7.43^ and thereby promote a clockwise rotation of TM7, disproportionately reducing arrestin recruitment. Moreover, we hypothesized that this effect would be larger for MP and DAMGO than LFT because the mutation will disrupt the former ligands’ hydrogen bonding interactions with Y326^7.43^, but not LFT’s π-π stacking interaction. In accord with these hypotheses, the Y326^7.43^F mutation eliminated detectable β-arrestin 2 recruitment for MP and reduced maximal β-arrestin 2 recruitment for DAMGO and LFT by 94% and 57%, respectively. The same mutation led to a lesser reduction in Gi activation, reducing maximal Gi1 activation for MP, DAMGO, and LFT by 68%, 36%, and 13%, respectively (Extended Data Fig. 7).

### Conclusion

The μOR, the endogenous enkephalin and β-endorphin receptor, is targeted by powerful pain medications with distinct scaffolds, such as morphine and fentanyl. Recent structures of the μOR bound to the morphinan antagonist βFNA, the morphinan agonist BU72 and the peptide agonist DAMGO, revealed how the orthosteric pocket can accommodate chemically diverse scaffolds. Here we provided high-resolution structural and mechanistic information for the partial-agonist MP and full-agonist LFT bound to the μOR in complex with Gi1. The hydrophobic nature of the orthosteric ligand binding site is responsible for the high-affinity of the predominantly hydrophobic LFT molecule, while a distinct sub-pocket is occupied solely by the MP scaffold. Molecular dynamics simulations provide novel insights into the plasticity of μOR activation and reveal how differences in ligand interactions within the orthosteric pocket can lead to different conformations of the intracellular G protein and β-arrestin coupling interface, resulting in marked differences in recruitment of β-arrestins and very distinct G protein activation profiles (Fig. 1b).

It has been proposed that the lack of respiratory depression in several G protein biased μOR agonists can be attributed to partial agonism for G protein activation^18^. However, the different G protein activation profiles may also have a critical role in these effects. It has previously been shown that MP is approximately ten-fold more potent than morphine and produces the same level of analgesia, yet MP is much less efficacious in promoting respiratory depression than morphine^11^. Notably, the G protein signaling profile of morphine is comparable to that of DAMGO and LFT, but very different from MP (Extended Data Fig. 11). In our assays, morphine displayed the highest potency (1.25 nM) and efficacy (106%) at Gz and the lowest at Gi2 (6.1 nM) with a potency rank order of Gz∼Gi1>GoB∼Gi3>GoA∼Gi2, while its efficacy at all Gα-subtypes was higher than MP. These results suggest that the G protein subtype(s) responsible for analgesia may be different from those primarily responsible for respiratory depression. Taken together, our findings emphasize the importance of a more complete functional characterization of μOR agonists for activation of all relevant G protein subtypes, as well as β-arrestin recruitment.

## Author Contributions

W.H. and H.W. prepared protein samples for structural studies and conducted biochemical assays. Q.Q. and A.B.S collected and processed the cryo-EM data and generated the maps. Q.Q., M.J.R. and A.B.S. built and refined models. S.C synthesized MP. T.C. and J.F.D. performed signaling profile assays under the supervision of B.L.R. A.I. performed NanoBiT experiments. D.A. and J.M.P. performed and analyzed molecular dynamics simulations under the supervision of R.O.D. Q.Q., W.H., D.A., J.M.P., A.B.S., S.M., R.O.D., B.K.K. and G.S. interpreted the data and wrote the manuscript with inputs from all authors. B.K.K. and G.S. supervised the project.

## Acknowledgements

This work was supported by the Swiss National Science Foundation Early Postdoctoral Mobility grant P2ELP3_187989 (D.A.), the European Molecular Biology Organization Long-Term Fellowship ALTF 544-2019 (D.A.), a Stanford Graduate Fellowship (J.M.P.), the National Institutes of Health grants R01GM127359 (R.O.D.), DA045884 (S.M.), R37DA036246 (B.K.K. and G.S.) and the Mathers Foundation (G.S. and B.K.K.). B.K.K. is a Chan Zuckerberg Biohub Investigator. This research used resources of the Oak Ridge Leadership Computing Facility, which is a U.S. Department of Energy Office of Science User Facility supported under contract DE-AC05-00OR22725. A.I. was funded by the PRIME 19gm5910013 (A.I.) and the LEAP 19gm0010004 and the BINDS JP20am0101095 from the Japan Agency for Medical Research and Development (AMED), KAKENHI 21H04791 and 21H05113 from the Japan Society for the Promotion of Science (JSPS), JST Moonshot Research and Development Program JPMJMS2023 from Japan Science and Technology Agency (JST). The authors thank Francois Marie Ngako Kadji, Kayo Sato, Yuko Sugamura and Ayumi Inoue at Tohoku University for plasmid construction and the cell-based GPCR assays; and Carl-Mikael Suomivuori, Yianni Laloudakis, Scott Hollingsworth and Naomi Latorraca for helpful discussions.

## Data and materials availability

The atomic coordinates for MP-μOR-Gi1 and LFT-μOR-Gi1-scFv complexes have been deposited in the Protein Data Bank with the accession codes 7T2G and 7T2H, respectively. The EM maps for MP-μOR-Gi1 and LFT-μOR-Gi1-scFv complexes have been deposited in EMDB with the accession codes EMD-25612 and EMD-25613, respectively. The composite non-model-based density modified map for MP-μOR-Gi1 is deposited to The Electron Microscopy Data Bank (EMDB) as the main map and used for model building. The locally refined individual maps are deposited as additional maps.

## Methods

### Expression and purification of μOR

#### A modified *M. musculus* (GenBank

AAB60673.1)μOR construct with removable N-terminal Flag-tag and C-terminal 10X histidine tag was used in this study. N-terminal residues (1-63) of μOR were replaced with the thermostabilized apocytochrome b_562_RIL from *Escherichia coli* (M7W, H102I and R106L) (BRIL) protein and a linker sequence (GSPGARSAS). N-terminal Flag-tag and C-terminal histidine tag were removable with rhinovirus 3C protease.μOR was expressed and purified as previously described^35^. Briefly, sf9 cells (Expression System) was infected with baculovirus at a density of 4×10^6^ cells/mL and incubated for 60 hours at 27 °C. Cell membrane was solubilized in n-dodecyl-b-D-maltoside (DDM, Anatrace) and 3-[(3-Cholamidopropyl)-dimethylammonio]-1-propanesulphonate (CHAPS, Anatrace) and purified by Ni-NTA resin (Thermo Scientific). The elute was further purified by M1 anti-Flag immunoaffinity resin and changed to a final buffer comprised of 25 mM HEPES pH 7.4, 100 mM NaCl, 0.01% lauryl maltose neopentyl glycol (L-MNG, Anatrace) and 0.001% cholesterol hemisuccinate (CHS, Sigma) by size exclusion chromatography. The peak fractions were collected and concentrated to ∼100μM.

### Assembly and purification of the LFT-μOR-Gi1-scFv16 and MP-μOR-Gi1 complexes

The heterotrimeric Gi1 (Gαi1/Gβ2/Gγ1) and scFv16 was expressed and purified as previously described^28^. Briefly, the heterotrimeric Gi was expressed in Trichuplusia ni Hi5 cells and purified in a final buffer containing 20 mM HEPES pH 7.4, 100 mM NaCl, 0.05% DDM, 1 mM MgCl_2_, 10 μM GDP and concentrated to ∼20 mg/mL for complexing. The scFv16 was expressed and secreted from Trichuplusia ni Hi5 and purified in a buffer containing 20 mM HEPES 7.4, 100 mM NaCl and was concentrated to ∼80 mg/mL for final use. Before complexing with μOR, the purified Gi heterotrimer was exchanged to L-MNG by adding equal volume of 20 mM HEPES 7.5, 50 mM NaCl, 1% L-MNG, 0.1% CHS, 1 mM MgCl_2_, 50 μM TCEP and 10 μM GDP at room temperature for 1 hour.

To prepare LFT bound μOR-Gi1 complex, the receptor was incubated with LFT at a final concentration of 1 mM at 4°C for 1 hour. Ligand-bound μOR was mixed with a 1.2 molar excess of heterotrimeric Gi1 and incubated at room temperature for 1 hour. Apyrase and λ-phosphatase (New England Biolabs) were added to the complex and incubated for another 1 hour at 4°C. The mixture was purified by M1 anti-Flag affinity chromatography to remove excess Gi1 protein and gradually change to a final buffer containing 20 mM HEPES 7.4, 100 mM NaCl, 0.0075% L-MNG, 0.001% CHS, 0.0025% glycol-diosgenin (GND, Anatrace), 250 nM lofentanil, 2 mM EDTA and 200 μg/mL Flag peptide. A 1.25 excess of scFv16 was added to the complex and incubated at room temperature for 1 hour. The LFT-μOR-Gi1_-_scFv16 complex was further purified by size exclusion chromatography on a Superdex 200 10/300 GL column (GE healthcare) in 20 mM HEPES 7.4, 100 mM NaCl, 0.00075% L-MNG, 0.0001% CHS, 0.00025% GDN and 250 nM LFT. Peak fractions were concentrated ∼20 mg/mL for electron microscopy studies.

For MP-μOR-Gi1 complex assembly, 500 μM MP was added to purified μOR while 1% L-MNG was added to purified Gi1. Both mixtures were incubated on ice for 1 h. After that, MP-bound μOR was mixed with a 1.5 molar excess of Gi1 heterotrimer and extra TCEP was added to maintain 100 μM TCEP concentration. The coupling reaction was allowed to proceed for another 1 h on ice, followed by addition of apyrase to catalyze GDP hydrolysis. The reaction mixture was left on ice overnight to allow stable complex formation. After that, the complexing mixture was purified by M1 anti-Flag affinity chromatography and eluted in 20 mM HEPES pH 7.5, 100 mM NaCl, 0.003% L-MNG, 0.001% glyco-diosgenin (GDN), 0.0004% CHS, 10 μM MP, 5 mM EDTA and Flag peptide. After elution, 100 μM TCEP was added to provide a reducing environment. Finally, the μOR–Gi1 complex was purified by size exclusion chromatography on a Superdex 200 10/300 gel filtration column in 20 mM HEPES pH 7.5, 100 mM NaCl, 5μM MP, 0.003% L-MNG and 0.001% GDN with 0.0004% CHS total. Peak fractions were concentrated to ∼10 mg/mL for electron microscopy studies.

### Cryo-EM sample preparation and image acquisition

The homogeneity of purified LFT-μOR-Gi1-scFv16 or MP-μOR-Gi1 complex was evaluated by negative stain EM^36^. For cryo-EM preparation of LFT-μOR-Gi1-scFv16 complex, 3.5 μL sample with addition of 0.05% β-OG was directly applied to glow-discharged 200 mesh gold grids (Quantifoil R1.2/1.3) and vitrified using a FEI Vitrobot Mark IV (Thermo Fisher Scientific). Images were collected on a Titan Krios (SLAC/Stanford) operated at 300 keV at a nominal magnification of 130,000X using a Gatan K2 Summit direct electron detector in counting mode, corresponding to a pixel size of 1.06Å. Movie stacks were obtained with a defocus range of -1.0 to -2.0 μm, using SerialEM^36^ with a set of customized scripts enabling automated low-dose image acquisition. Each movie stack was recorded for a total of 8 seconds with 0.2s exposure per frame and exposure dose set to 7 electrons per pixel per second.

MP-μOR-Gi1 protein complex was vitrified in a manner similar to LFT-μOR-Gi1-scFv16 complex, except on a 300 mesh UltrAuFoil grid (Quantifoil R1.2/1.3) and imaged at a magnification of 165,000 (0.82 Å/pixel). Movie stacks with dose fractioned over 40 frames, were recorded with a dose rate of 1.4 e/Å^2^ (6.27 e/pixel/second) using counting mode with a defocus range of -0.8 ∼-1.8 μm for MP-μOR-Gi protein complex using SerialEM.

### Cryo-EM data processing

Datasets for LFT-μOR-Gi1-scFv16 and MP-μOR-Gi1 complex was processed using CryoSPARC (v3.2)^37^ and Relion (v3.1)^38^ respectively. For both LFT-μOR-Gi1-scFv16 and MP-μOR-Gi1 complex, a total 1853 or 1931 image stacks were subjected to beam-induced motion correction using CryoSPARC patch motion correction algorithm and MotionCor2^39^, respectively (Extended Data Figs. 2-3). Contrast transfer function parameters for each micrograph were estimated from the exposure-weighted averages of all frames by CryoSPARC patch CTF algorithm and Gctf (v1.06)^40^, implemented in Relion. Particles were autopicked using reference-based picking, extracted with a box size of 256 pixels, and subjected to several rounds of 2D classification to remove contaminants. Initial maps were generated using stochastic gradient descent-based *ab-initio* refinement in CryoSPARC and Relion. Selected particle sets were further cleaned with several rounds of 3D classification. The final dataset of 152,809 particles for LFT-μOR-Gi1-scFv16 was subjected to 3D non-uniform refinement^41^ after Ctf refinement, generating a 3.2Å map sharpened with CryoSPARC. The final dataset of 413,821 particles for MP-μOR-Gi1 was subjected to 3D auto-refinement after Bayesian polishing and Ctf refinement, which generated a 2.5Å map. Resolution of these maps were estimated internally in CryoSPARC and Relion by gold standard Fourier shell correlation using the 0.143 criterion. The MP-μOR-Gi1 map was further locally refined with finer angular sampling (0.9 degrees) using masks including only the receptor or G protein heterotrimer in Relion. Locally refined MP-μOR-Gi1 maps were density modified and sharpened with Resolve Cryo-EM^42^ procedure in Phenix (dev-4271) using non-model-based algorithms, which yielded improved local maps with better than 2 Åresolution in stable areas (Extended Data Figs. 2-3). Composite maps for MP-μOR-Gi1 were generated using ChimeraX (v1.2)^43^. The composite non-model-based density modified map for MP-μOR-Gi1 is deposited to The Electron Microscopy Data Bank (EMDB) as the main map. The locally refined maps are deposited as additional maps. Local resolution estimation was performed with CryoSPARC’s and Relion’s own local resolution estimation algorithms using half maps.

### Model building and refinement

Initial ligand models were generated by the Edit tool implemented in Phenix^44^, using LFT or MP SMILES. Together with individual protein chains from the DAMGO-μOR-Gi1-scFv16 structure, all models were rigid-body docked into the corresponding cryo-EM density map in Chimera^45^, followed by iterative manual adjustment in COOT^46^ and real space refinement in Phenix. Ligand coordination was further optimized by GemSpot^47^. The model statistics were validated using Molprobity^48^. Structural figures were prepared in Chimera (v1.15), ChimeraX (v1.2) or PyMOL (Schrödinger) (The PyMOL Molecular Graphics System, Version 2.0 Schrödinger, LLC.) (https://pymol.org/2/). The final refinement statistics are provided in Extended Data Table 2.

### Generating fentanyl and carfentanil docking models

μOR was prepared for docking by the addition of missing sidechain atoms and hydrogen bonding optimization with Schrödinger’s Maestro protein preparation wizard (Schrödinger Release 2018-4: Maestro, Schrödinger, LLC, New York, NY, 2018). Glide extra precision (XP) docking^49^ was executed on the prepared structure with fentanyl or carfentanil. Docking identified largely identical poses for carfentanil but two distinct high-scoring poses for fentanyl.

### System setup for molecular dynamics simulations

We performed simulations of μOR bound to MP, LFT, DAMGO, and fentanyl. We initiated MP-and LFT-bound simulations from the MP-μOR-Gi1, and LFT-μOR-Gi1-scFv16 structures reported in this manuscript, respectively. We initiated the DAMGO-bound simulations from the previously published DAMGO-μOR-Gi1-scFv16 structure (PDB ID: 6DDF)^28^. We initiated fentanyl-bound simulations from the LFT-μOR-Gi1-scFv16 structure reported in this manuscript and converted LFT to fentanyl using Maestro (Schrödinger Release 2018-4: Maestro, Schrödinger, LLC, New York, NY, 2018). We also initiated fentanyl-bound simulations from the top two unique docking poses of fentanyl. In all simulation conditions, we removed the Gi1 and scFv16 chains. For MP, LFT, and DAMGO simulations, we performed six independent simulations, each 3.5μs in length. For fentanyl simulations, we performed ten independent simulations, each approximately 1 μs in length. For each simulation, initial atom velocities were assigned randomly and independently.

For all simulation conditions, the protein structures were aligned to the Orientations of Proteins in Membranes^50^ entry for 5C1M (active μOR bound to BU72^31^) using PyMOL, and crystal waters from 5C1M were incorporated. Prime (Schrödinger)^51^ was used to model missing side chains, and to add capping groups to protein chain termini. Parameters for MP, LFT and fentanyl were generated using the Paramchem webserver^52-54^. Parameters for DAMGO were obtained as previously described^30^. Protonation states of all titratable residues were assigned at pH 7, except for D^2.50^ and D^3.49^, which were protonated (neutral) in all simulations, as these conserved residues are reported to be protonated in active-state GPCRs^55,56^. Histidine residues were modeled as neutral, with a hydrogen atom bound to either the delta or epsilon nitrogen depending on which tautomeric state optimized the local hydrogen-bonding network. Using Dabble (R. Betz, Dabble (v2.6.3), Zenodo (2017); doi:10.5281/zenodo.836914.), the prepared protein structures were inserted into a pre-equilibrated palmitoyl-oleoyl-phosphatidylcholine (POPC) bilayer, the system was solvated, and sodium and chloride ions were added to neutralize the system and to obtain a final concentration of 150 mM. The final systems comprised approximately 59,000 atoms, and system dimensions were approximately 80×80×90 Å.

### Molecular dynamics simulation and analysis protocols

We used the CHARMM36m force field for proteins, the CHARMM36 force field for lipids and ions, and the TIP3P model for waters^57-59^. All simulations were performed using the Compute Unified Device Architecture (CUDA) version of particle-mesh Ewald molecular dynamics (PMEMD) in AMBER18^60^ on graphics processing units (GPUs).

Systems were first minimized using three rounds of minimization, each consisting of 500 cycles of steepest descent followed by 500 cycles of conjugate gradient optimization. 10.0 and 5.0 kcal.mol^−1^·Å^−2^ harmonic restraints were applied to protein, lipids, and ligand for the first and second rounds of minimization, respectively. 1 kcal.mol^−1^·Å^−2^ harmonic restraints were applied to protein and ligand for the third round of minimization. Systems were then heated from 0 K to 100 K in the NVT ensemble over 12.5 ps and then from 100 K to 310 K in the NPT ensemble over 125 ps, using 10.0 kcal.mol^−1^·Å^−2^ harmonic restraints applied to protein and ligand heavy atoms.

Subsequently, systems were equilibrated at 310 K and 1 bar in the NPT ensemble, with harmonic restraints on the protein and ligand non-hydrogen atoms tapered off by 1.0 kcal.mol^-1^·Å^-2^ starting at 5.0 kcal.mol^-1^·Å^-2^ in a stepwise fashion every 2 ns for 10 ns, and then by 0.1 kcal.mol^-1^·Å^-2^ every 2 ns for 20 ns. Production simulations were performed without restraints at 310 K and 1 bar in the NPT ensemble using the Langevin thermostat and the Monte Carlo barostat, and using a timestep of 4.0 fs with hydrogen mass repartitioning^61^. Bond lengths were constrained using the SHAKE algorithm^62^. Non-bonded interactions were cut off at 9.0 Å, and long-range electrostatic interactions were calculated using the particle-mesh Ewald (PME) method with an Ewald coefficient of approximately 0.31 Å, and 4th order B-splines. The PME grid size was chosen such that the width of a grid cell was approximately 1 Å. Trajectory frames were saved every 200 ps during the production simulations.

The AmberTools17 CPPTRAJ package was used to reimage trajectories^63^. Simulations were visualized and analyzed using Visual Molecular Dynamics (VMD)^64^ and PyMOL.

To construct the probability distributions shown in Figs. 4b and 5d, we used trajectory frames from all simulations under each condition and applied a Gaussian kernel density estimator. In Fig. 4b, the distances were measured between (i) the side-chain oxygen of Y128^2.64^ and the closer of the side-chain oxygen and nitrogen of Q124^2.60^, and (ii) the side-chain oxygen of Y326^7.43^ and the closer of the side-chain oxygen and nitrogen of Q124^2.60^. In Fig. 5d, the distance was measured between the Cα atoms of P333^7.50^ and L110^2.46^.

### NanoBiT-β-arrestin recruitment assay

μOR-induced β-arrestin recruitment was measured by a NanoBiT-β-arrestin recruitment assay, in which interaction between μOR and β-arrestin was monitored by a NanoBiT enzyme complementation system^65^. Human full-length β-arrestin1 and β-arrestin2 were N-terminally fused to a large fragment (LgBiT) of the NanoBiT luciferase with a 15-amino acid (GGSGGGGSGGSSSGG) flexible linker (Lg-βarr1 and Lg-βarr2, respectively). Human full-length μOR (WT or mutants) was C-terminally fused to a small fragment (SmBiT) with the 15-amino acid flexible linker (μOR-Sm). Lg-βarr1, Lg-βarr2 and μOR-Sm constructs were inserted into a pCAGGS expression plasmid vector. HEK293A cells (Thermo Fisher Scientific) were seeded in a 10-cm culture dish at a concentration of 2 × 10^5^ cells mL^-1^ (10 mL per dish in DMEM (Nissui Pharmaceutical) supplemented with 10% fetal bovine serum (Gibco), glutamine, penicillin and streptomycin) 1-day before transfection. Transfection solution was prepared by combining 20 μL (per dish, hereafter) of polyethylenimine solution (Polysciences; 1 mg/mL) and a plasmid mixture consisting of 500 ng Lg-βarr1 or Lg-βarr2 and 1μg μOR-Sm in 1 mL of Opti-MEM (Thermo Fisher Scientific). After incubation for 1-day, transfected cells were harvested with 0.5 mM EDTA-containing Dulbecco’s PBS, centrifuged and suspended in 10 mL of HBSS containing 0.01% bovine serum albumin (BSA; fatty acid–free grade; SERVA) and 5 mM HEPES (pH 7.4) (assay buffer). The cell suspension was dispensed in a white 96-well culture plate at a volume of 80 μL per well and loaded with 20 μL of 50 μM coelenterazine (Carbosynth; final concentration at 10 μM) diluted in the assay buffer. After 2-h incubation at room temperature, the plate was measured for baseline luminescence (Spectramax L, Molecular Devices) and 20 μL of 6X test compounds diluted in the assay buffer or the assay buffer alone (vehicle) were manually added. The plate was read for 15 min with an interval of 20 sec at room temperature. Luminescence counts recorded from 10 min to 15 min after compound addition were averaged and normalized to the initial counts. The fold-change signals were further normalized to the vehicle-treated signal and were plotted as a G protein dissociation response. The resulting luminescent counts were fitted to a four-parameter sigmoidal concentration-response curve using Prism 8 software (GraphPad Prism) and pEC_50_ values (negative logarithmic values of EC_50_ values) and *E*_max_ values (“Top” – “Bottom”) were obtained from the curve fitting and used for calculation of mean and SEM. For individual experiments, we calculated *E*_max_/EC_50_ of LFT or MP relative to that of DAMGO, a dimensionless parameter known as relative intrinsic activity (RAi)^66^ to indicate agonist activity, and used its base-10 log-transformed value (Log RAi) to obtain mean and SEM.

### NanoBiT-G protein dissociation assay

μOR-induced Gi1 activation was measured by a NanoBiT-G protein dissociation assay^67^, in which dissociation of Gαi1 subunit from Gβ1γ2 subunit was monitored by the NanoBiT system. Specifically, a NanoBiT-Gi1 protein consisting of LgBiT-containing Gα_i1_ subunit (Lg-Gαi1), SmBiT-fused Gγ_2_ subunit harboring a C68S mutation (Sm-Gγ2 (C68S)) and untagged Gβ1 subunit was expressed in HEK293A cells together with a μOR construct harboring N-terminal HA signal sequence and FLAG-epitope tag and C-terminal GFP (FLAG-μOR-GFP). Cell seeding and transfection were performed in the same procedures as described in the NanoBiT-β-arrestin recruitment assay except for a plasmid mixture (500 ng Lg-Gαi1, 2.5μg Gβ1, 2.5μg Sm-Gγ2 (C68S) and 1μg FLAG-μOR-GFP per 10-cm culture dish). After baseline luminescent measurement and addition of 20 μL test compounds, the plate was immediately placed in the luminescent microplate reader and measured for 5 min. Change in luminescent count from 3 min to 5 min was averaged and used to plot G-protein dissociation response. The G-protein dissociation signals were fitted to a four-parameter sigmoidal concentration-response curve and the Log RAi values were obtained as described above.

### Bioluminescence resonance energy transfer (BRET) assays

To measure μOR-mediated G protein dissociation of Gi1-, Gi2-, Gi3-, GoA-, GoB-, and Gz-containing heterotrimeric G proteins, HEK293T cells were co-transfected in a 1:1:1:1 ratio with human μOR and the optimal Gα-RLuc8, Gβ, and Gγ-GFP2 subunits described in the TRUPATH paper^24^. TransIT-2020 (Mirus Bio LLC) was used to complex the DNA at a ratio of 3 μL Transit per μg DNA, in Opti-MEM (Gibco-ThermoFisher) at a concentration of 10 ng DNA per μL Opti-MEM. After 16 hours, transfected cells were plated in poly-lysine coated 96-well white clear bottom cell culture plates in plating media (DMEM + 1% dialyzed FBS) at a density of 40-50,000 cells in 100 μL per well and incubated overnight. The next day, media was vacuum aspirated and cells were washed twice with 60 μL of assay buffer (20 mM HEPES, 1X HBSS, pH 7.4). Next, 60 μL of the RLuc substrate, coelenterazine 400a (Nanolight Technologies, 5 μM final concentration in assay buffer) was added per well, and incubated for 5 minutes to allow for substrate diffusion. Afterwards, 30 μL of drug (3X) in drug buffer (20 mM HEPES, 1X HBSS, 0.3% BSA, pH 7.4) was added per well and incubated for another 5 minutes. Plates were immediately read for both luminescence at 395nm and fluorescent GFP2 emission at 510 nm for 1 second per well using a Mithras LB940 multimode microplate reader. BRET ratios were computed as the ratio of the GFP2 emission to RLuc8 emission. Data were analyzed by a three-parameter nonlinear regression equation using Graphpad Prism 8 (Graphpad Software Inc., San Diego, CA). All experiments were repeated in at least three independent trials each with duplicate determinations.

To measure μOR recruitment of β-arrestin 1 and 2 subtypes, HEK293T cells were co-transfected in a 1:1:5 ratio with human μOR containing a C-terminal Renilla luciferase (RLuc8), human GRK2, and human β-arrestin1 or 2 containing an N-terminal mVenus. Each plate also contained wells co-transfected with pcDNA instead of mVenus-arrestin to measure background fluorescence. This assay was performed identically to the TRUPATH assay, except that the substrate used was coelenterazine h (Promega, 5μM final concentration in drug buffer) and plates were read for luminescence at 485 nm and fluorescent mVenus emission at 530 nm. BRET ratios were computed as the ratio of the mVenus to RLuc8 emission, and the net BRET was calculated by background subtraction of the BRET ratio from pcDNA-transfected wells. Data were analyzed by a three-parameter nonlinear regression equation using Graphpad Prism 8. All experiments were repeated in at least three independent trials each with duplicate determinations

