## Extended Data File for "Structural insights into distinct signaling profiles of the μOR activated by diverse agonists"

### Supplementary figure legends

**Extended Data Fig.1 | Activity characterization of diverse ligands on μOR.** **a,** Efficacy of compounds LFT, MP and Morphine at human μOR of Gi1, GoA, Go and Gz activation, and recruitment of β-arrestin-2 using the BRET assay are shown as a percentage of receptor activation relative to the full agonist, DAMGO.MP had significantly lower G-potein as well as arrestin among ligands tested (****p<0.0001). Statistical significance was determined using one-way ANOVA followed by Dunnett's multiple comparison test. **b,** Dose-dependent activation of Gi1 signaling in NanoBiT Gi1-dissociation assay and activation of β-arr1 and β-arr2 signaling in NanoBiT Arrestin-recruitment assay on wild type human μOR. Data are the means(+/- SE) from four independent experiments and summarized at bottom panel.

**Extended Data Fig.2 | Cryo-EM process for LFT-μOR-Gi1-scFv and MP-μOR-Gi1 complexes.** Representative raw micrographs (**a**) and 2D classification averages (**b**) for LFT-μOR-Gi1-scFv and MP-μOR-Gi1, respectively. **c,** Workflow of cryo-EM data processing of LFT (left) and MP (right). **d,** Angular distribution of projections for cryo-EM maps and gold-standard FSC curves of half-maps (0.143 cutoff). **e,** Local resolution of LFT-μOR-Gi1-scFv cryo-EM map. **f**, Local resolution of composite cryo-EM maps of MP-μOR-Gi1 after local refinement and non-model based density modification (Phenix Resolve Cryo-EM).

**Extended Data Fig.3 | Representative regions of high-quality cryo-EM density and corresponding models.** Exhibition of transmembrane domains (TM) for (**a**) LFT-μOR-Gi1-scFv complex and (**b**) MP-μOR-Gi1 complex. and (**c**) Gα subunit αN and α5 helices, one CHS molecule within close proximity to TM domain, MP ligand and four water molecules distributed along the central funnel of μOR (shown as wire and red dot) which overlay quite well with waters identified in 2.0Å agonist BU72-μOR crystal structure (blue dot) (PDB: 5c1m). **d, e,** Representative density views for TM3 (**d**) and Gβ residues (**e**) of MP-μOR-Gi1 complex map via non-model-based density modification by Phenix Resolve Cryo-EM.

**Extended Data Fig.4 | Conserved μOR-Gi1 conformation activated by diverse agonists. a,** Alignment of MP (orange), LFT (red) and DAMGO (blue) bound μOR-Gi1 complexes onto BU72-bound μOR (grey), with nanobody and scFv removed for clarity. **b,** Close view of the ligand binding orthosteric pocket show nearly identical poses for residues involved in different agonists interaction, except that Q214 orients its side-chain towards TM3 upon MP engagement, which results in the loss of Q214-Y326 interaction observed among LFT, DAMGO and BU72. Orientation of GPCR activation feature motifs $P^{5.50}$-$I^{3.40}$-$F^{6.44}$ (**c**), $DR^{3.50}Y$ and $NP^{7.50}xxY^{7.53}$ (**d**), and residues lined on the major interface between μOR intracellular site composed of TM2-3, TM5-7 and ICL1-3 (**e**) and C-terminal α5 helix of Gα subunit (**f**) are quite similar, suggesting a canonical conformation for Gi1 heterotrimer coupled μOR activated by DAMGO, LFT and MP.

**Extended Data Fig.5 | Interaction network comparison among diverse μOR ligands. a,** Structurally distinct agonists such as morphinan BU72 (grey), enkephalin-like DAMGO (blue), synthetic LFT (magenta), novel alkaloid MP (green) and morphinan antagonist βFNA (black) all occupy the central pocket (cp) of wide orthostric binding site. **b**, Viewed from extracellular side, functional moieties of the BU72, DAMGO and LFT penetrate into a sub-pocket between TM2 and TM3 (sp1), while MP indole ring explores a new arena composed of TM1, TM2 and TM7 (sp2). **c,** Schematic interaction diagrams highlight a conserved salt-bridge/hydrogen-bond interaction between μOR D147 (red sphere) and a tertiary amine ($NH^+$) on BU72, DAMGO, LFT and MP, in addition to the major hydrophobic interaction network, calculated by Maestro (citation).

**Extended Data Fig.6 | Molecular basis for high potency of fentanyl analogues revealed by docking and simulation. a,** 2D diagram of fentanyl derivatives carfentanil and lofentanil. The 4-carbomethoxy moiety added on piperidinyl group of fentanyl confers carfentanil over 100 times more potent on μOR receptor, while further addition of 3-methyl group slightly enhances lofentanil in efficacy compared to carfentanil. **b-d,** Docking poses of fentanyl and carfentanil based on LFT coordinate within LFT-μOR cryo-EM structure. LFT (magenta stick) is overlaid with (**b**) fentanyl docking pose 1 (orange stick), (**c**) fentanyl docking pose 2 (yellow stick) and (**d**) carfentanil (teal stick). **e**, Simulations with fentanyl in three different binding poses: (i) fentanyl modelled by modifying LFT in the LFT cryo-EM structure (“the cryo-EM pose”), (ii) fentanyl docking pose 1, which has a GlideScore and Emodel better than fentanyl docking pose 2, (iii) fentanyl docking pose 2, which is a pose more similar to the LFT cryo-EM pose. For each simulation, the RMSD of fentanyl is computed after alignment on the receptor.

**Extended Data Fig.7 | Signaling characterization of μOR mutants upon various ligands. a**, NanoBiT based G-protein dissociation assay and β-arr2 recruitment assay of wild type human μOR and mutants(I322A,Q124A,Y326F, W293F) on DAMGO, MP and LFT. **b**, and their summary. Data are the means (+/- SE) from four independent experiments.

**Extended Data Fig. 8 | Comparison between binding pocket conformations observed in cryo-EM structures and in simulation.** In simulation, stable π-π stacking between LFT and $Y326^{7.43}$, together with the strong $Y326^{7.43}$–$Q124^{2.60}$ and $Q124^{2.60}$– $Y128^{2.64}$ interactions, favors the inwards cryo-EM conformation of $Y326^{7.43}$ (right panel). MP sterically blocks the $Y326^{7.43}$–$Q124^{2.60}$ interaction. In addition, the hydrogen bond between $Y326^{7.43}$ and MP observed in the cryo-EM structure is not stable in simulation, as it can be replaced by a water-mediated interaction (left panel). Even though the hydrogen bond between $Y326^{7.43}$ and DAMGO can also be replaced by a water-mediated interaction, the $Y326^{7.43}$–$Q124^{2.60}$ interaction, which is often formed in simulation in the presence of DAMGO, results in an intermediate positioning of $Y326^{7.43}$ (middle panel).

**Extended Data Fig. 9 | The transition from the canonical active conformation to the alternative conformation involves changes in an interaction network in the core of the receptor.** The transition from the canonical active conformation to the alternative conformation involves a shift in the hydrogen bonding network in the sodium binding pocket, as shown through representative frames from our μOR simulations. In the canonical active conformation, $N86^{1.50}$ forms a hydrogen bond with $S329^{7.46}$, and $D114^{2.50}$ forms a hydrogen bond with $N332^{7.49}$. In the alternative conformation, these interactions are broken and replaced by $D114^{2.50}$–$S329^{7.46}$ and $D114^{2.50}$–$N150^{3.35}$ hydrogen bonds. The μOR cryo-EM structure (grey) is represented by the MP-μOR-$G_i$ structure reported in this manuscript.

**Extended Data Fig. 10 | Comparison of the intracellular conformations observed for the μOR and the $AT_1R$. a,** Simulations indicate that in both the μOR and the $AT_1R$, the transition from the canonical active conformation to the alternative conformation involves a counterclockwise twist at TM7 (bottom panels), leading to relaxation of the kink in the NPxxY region and the inward movement of $P^{7.50}$ (top panels). Both at the μOR and the $AT_1R$, $P^{7.50}$ is translated inward in the alternative conformation with respect to the canonical active conformation and the inactive conformation (top panels). **b,** The μOR alternative conformation differs from the $AT_1R$ alternative conformation in that the intracellular end of TM7 shows an inward displacement in μOR compared to the $AT_1R$ alternative conformation. The interaction between $R^{3.50}$ and $D^{8.47}$ does not allow the downward $Y^{7.53}$ rotamer observed at the $AT_1R$. The canonical active and alternative conformations shown here are representative frames from our μOR simulations, and the inactive μOR and $AT_1R$ structures are the inactive μOR and $AT_1R$ crystal structures, respectively (PDB IDs: 4DKL and 4YAY).

**Extended Data Fig. 11 | Signaling profiles of μOR modulated by morphine.** Dose-dependent activation of Gi1, Gi2, Gi3, GoA, Go and Gz, and recruitment of β-arrestin-1 and β-arrestin-2 by morphine using the BRET assay.

**Extended Data Table 1. Summary of BRET assay results.** Data for all functional assays that were carried out in hMOR were normalized to Emax of DAMGO. The dose response curves were fit using a three-parameter logistic equation in GraphPad Prism, and the data are presented as mean EC50(pEC50 ± SEM) for assays run in duplicates at least three times.

**Extended Data Table 2. Cryo-EM data collection, refinement, and validation statistics.**

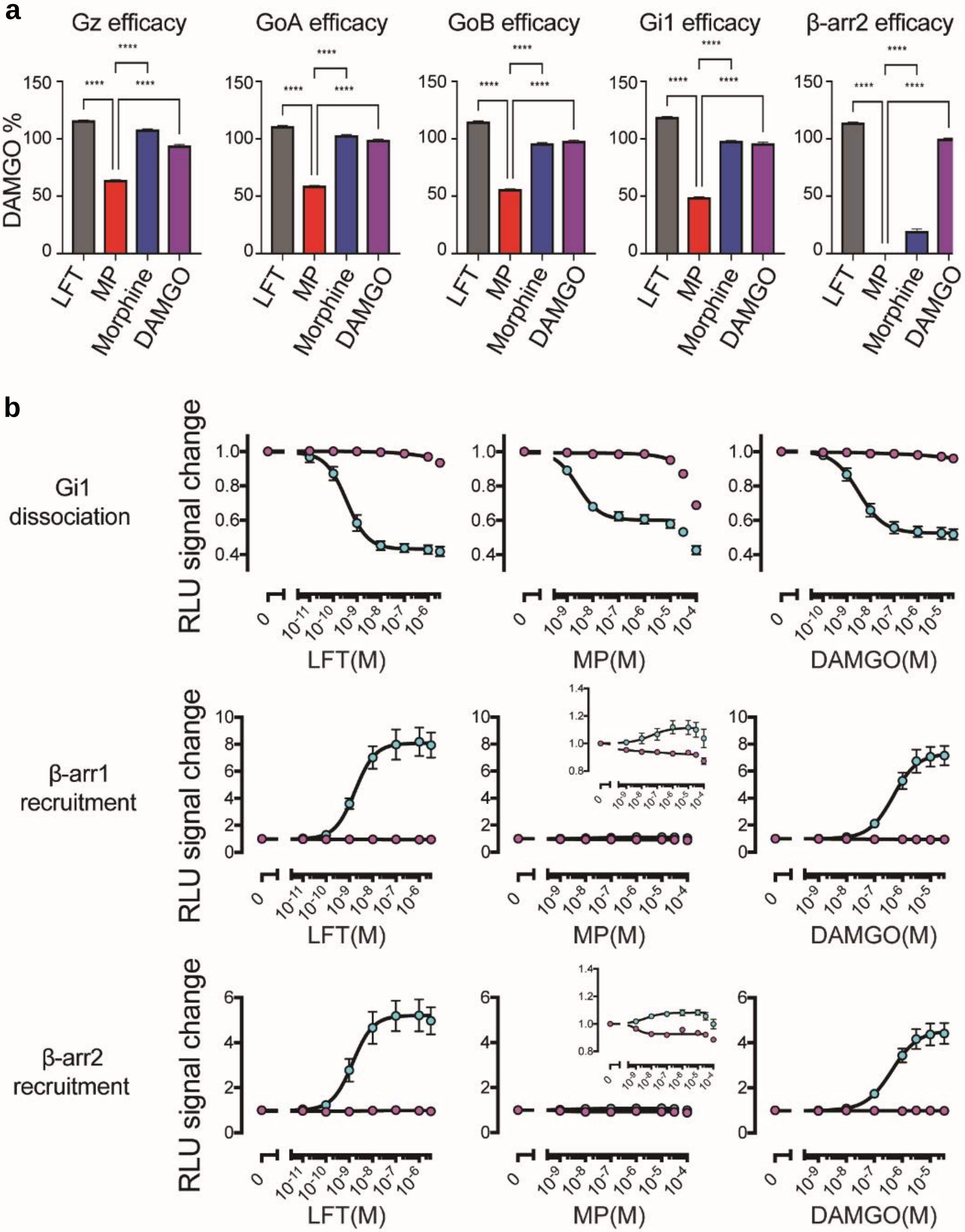


| | LFT | MP | DAMGO |
|---|---|---|---|
| Gi1 dissociation(pEC50) | 9.45±0.19 | 8.60±0.07 | 8.42±0.11 |
| β-arr1 recruitment(pEC50) | 8.79±0.10 | 7.27±0.37 | 6.33±0.05 |
| β-arr2 recruitment(pEC50) | 8.84±0.12 | 8.87±0.55 | 6.40±0.09 |

**Extended Data Fig.1 | Activity characterization of diverse ligands on μOR**

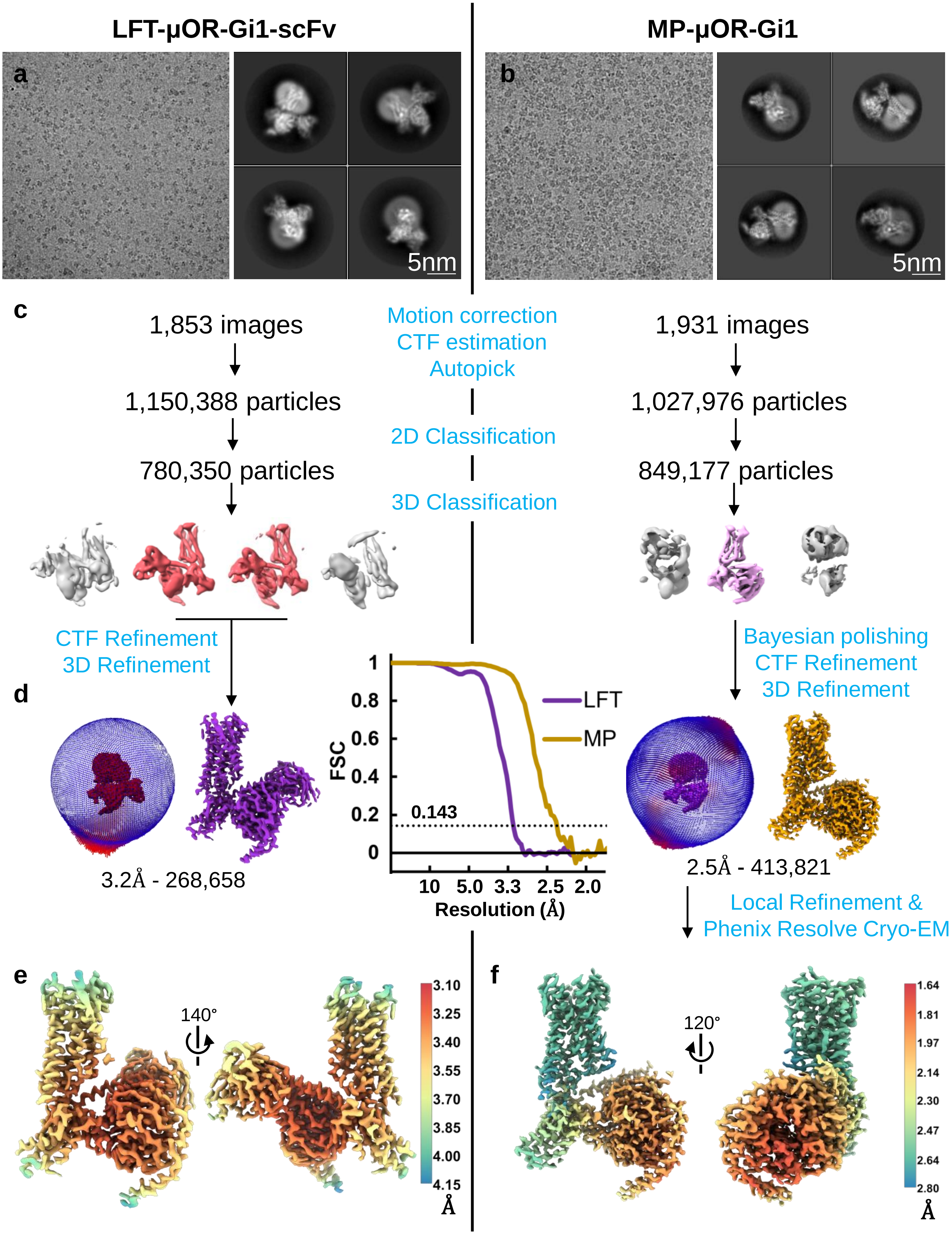


**Extended Data Fig.2 | Cryo-EM process for LFT-μOR-Gi1-scFv and MP-μOR-Gi1 complexes.**

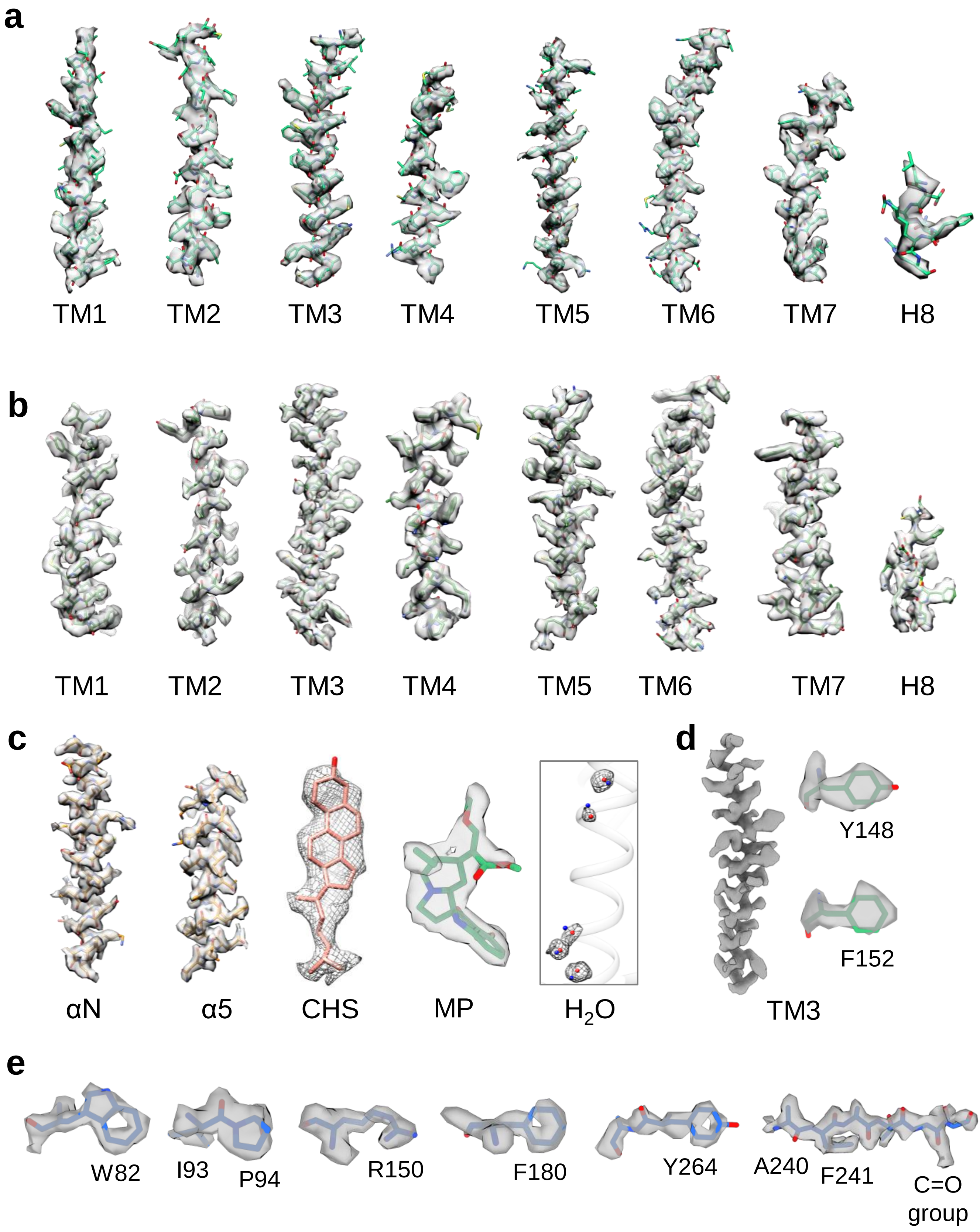


**Extended Data Fig.3 | Representative regions of high-quality cryo-EM density and corresponding models.**

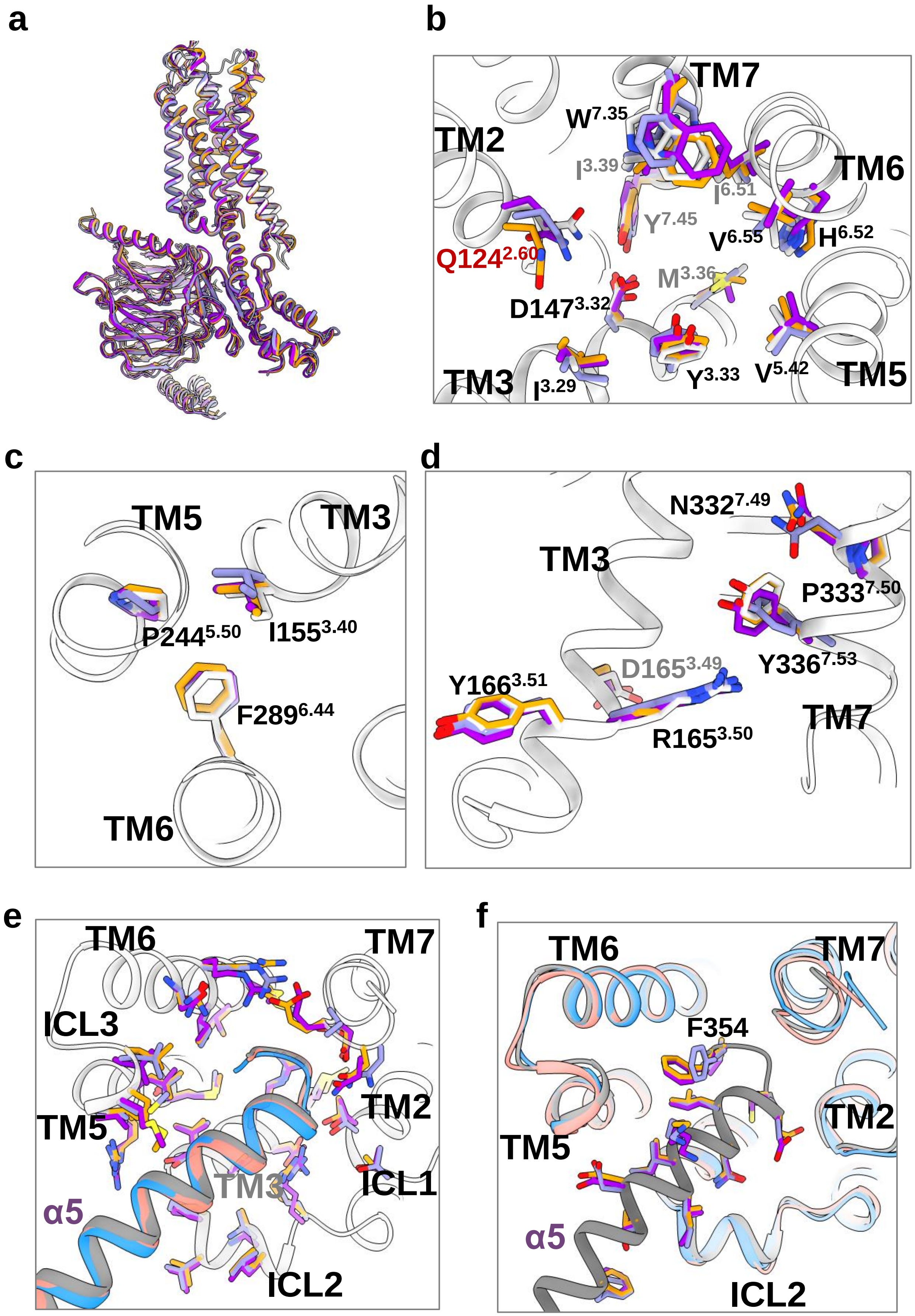


**Extended Data Fig.4 | Conserved μOR-Gi1 conformation activated by diverse agonists.**

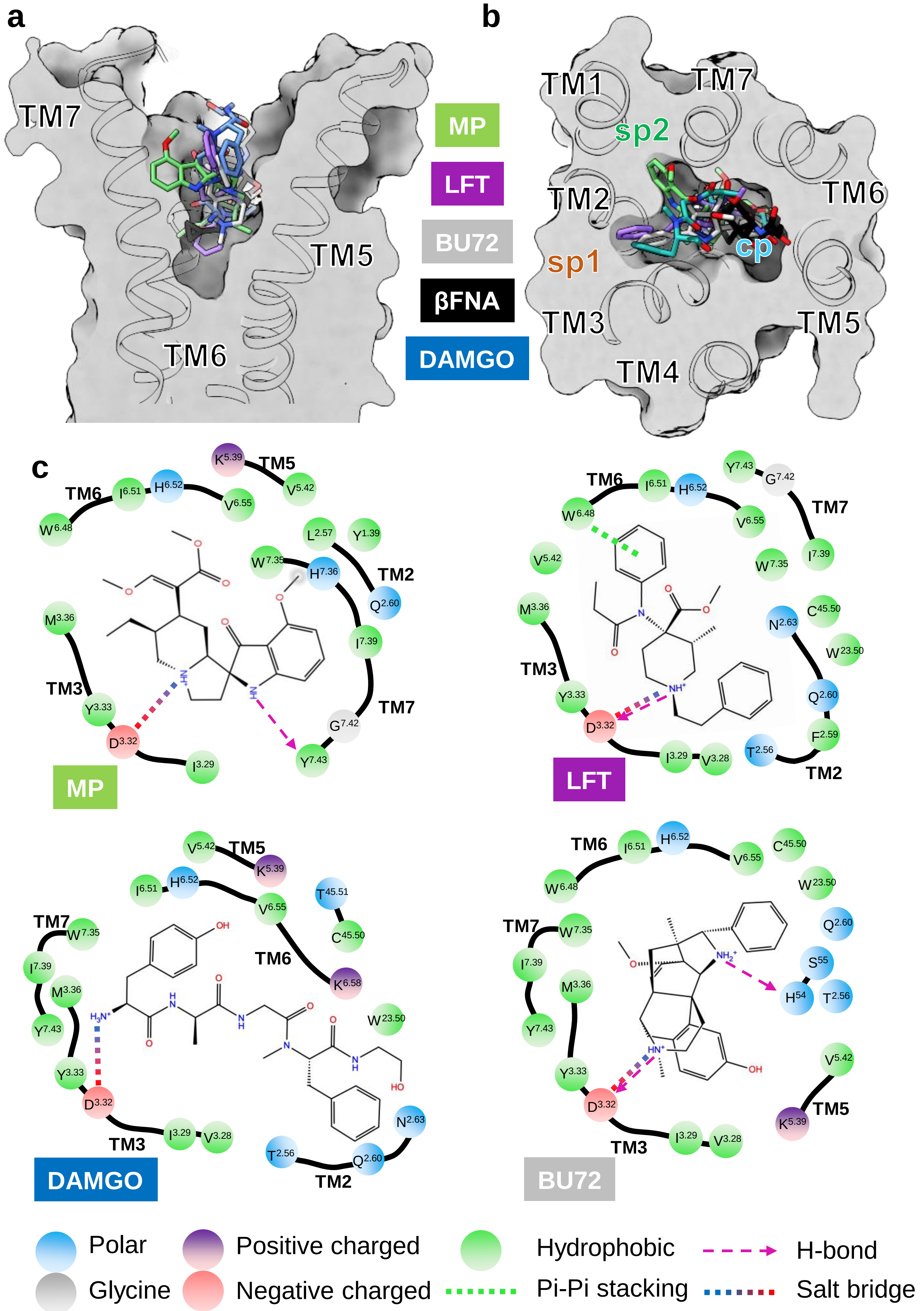


**Extended Data Fig.5 | Interaction network comparison among diverse μOR ligands.**

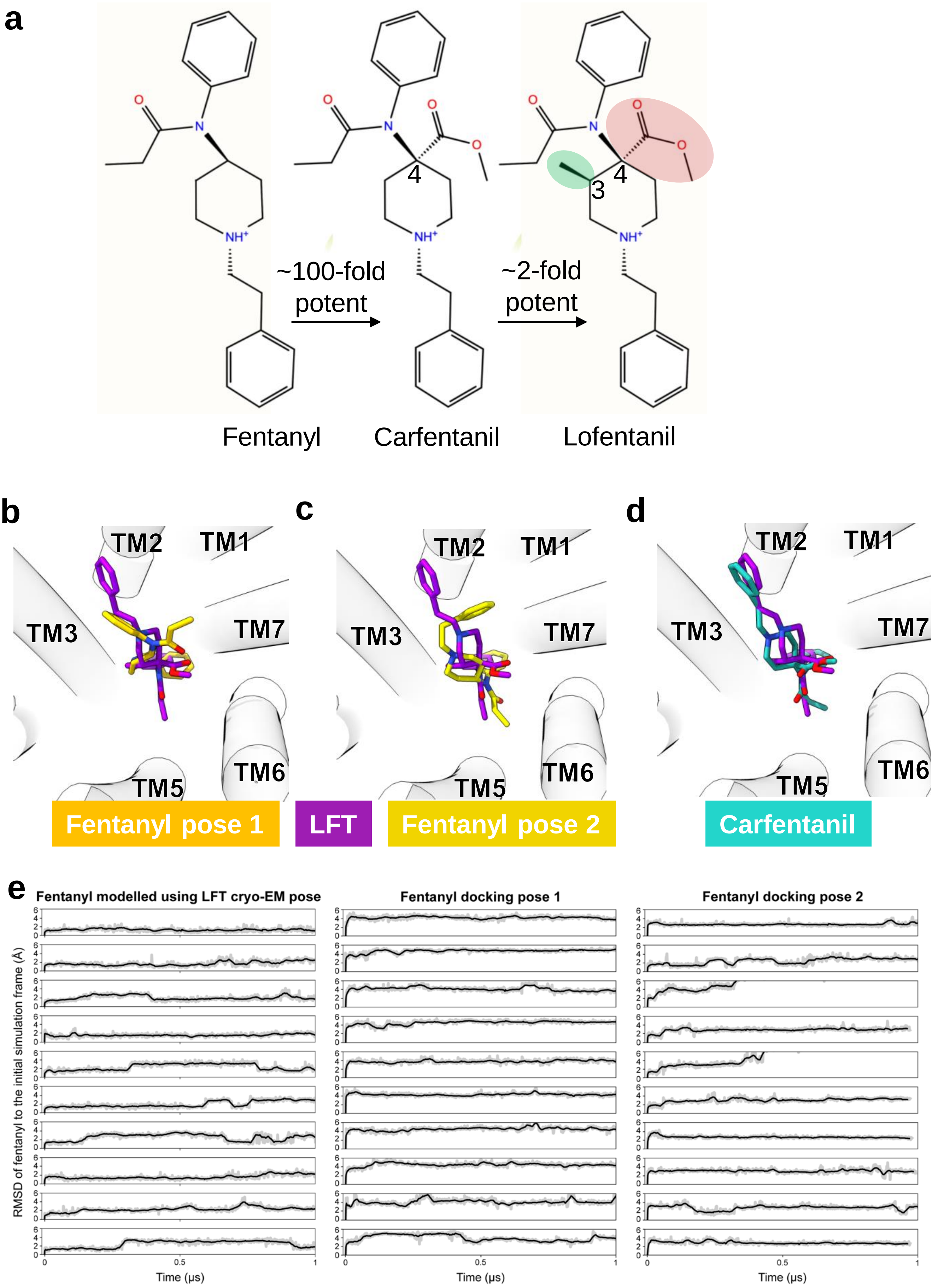


**Extended Data Fig.6 | Molecular basis for high potency of fentanyl analogues revealed by docking and simulation.**

a

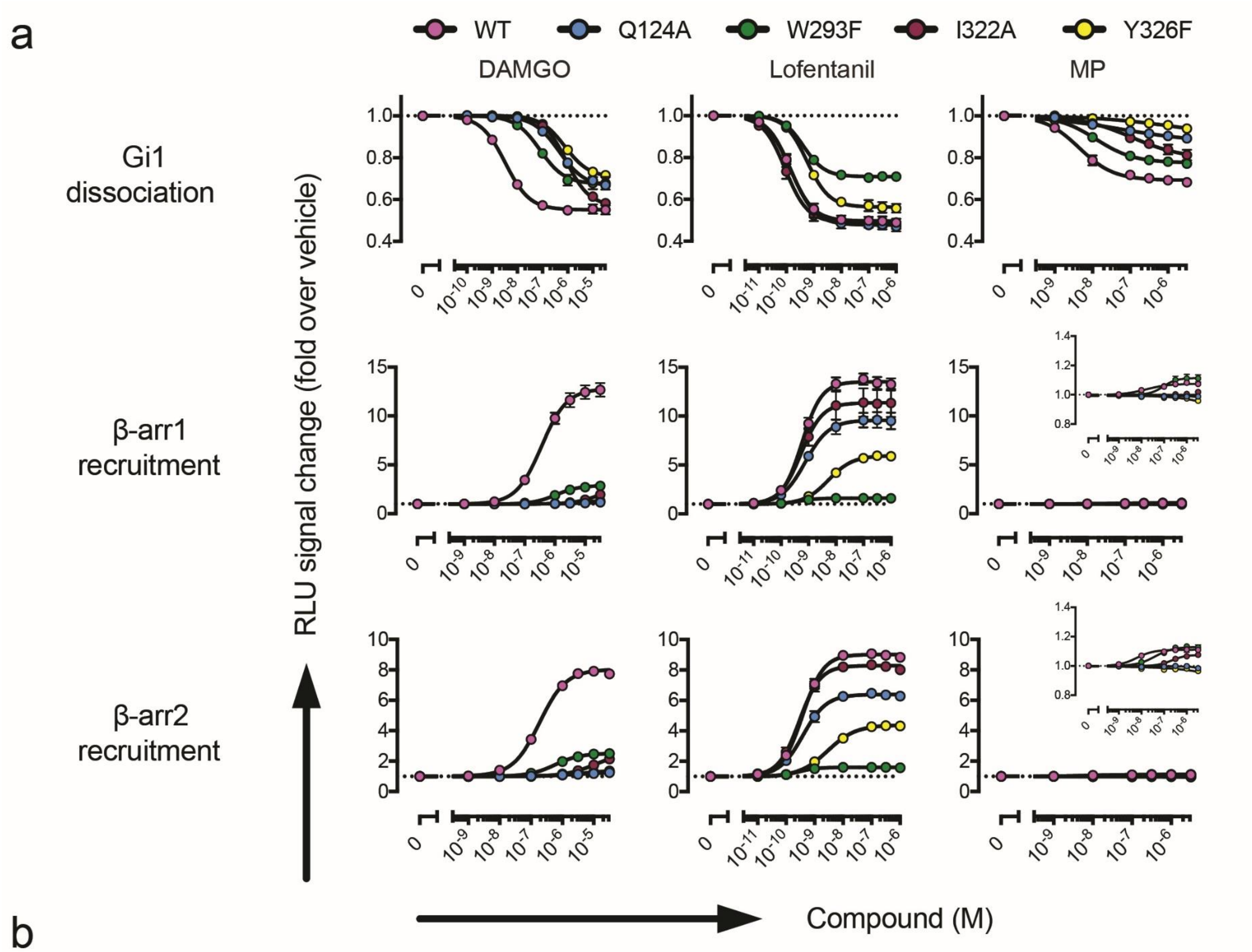


b

| DAMGO | WT | | Y326F | | I322A | | Q124A | | W293F | |
|---|---|---|---|---|---|---|---|---|---|---|
| | Emax% | pEC50 | Emax% | pEC50 | Emax% | pEC50 | Emax% | pEC50 | Emax% | pEC50 |
| Gi1 | 100% | 8.49 ± 0.05 | 65.5 ± 2.4 | 6.11 ± 0.04 | 96.6 ± 3.6 | 6.06 ± 0.03 | 76.5 ± 4.2 | 6.34 ± 0.03 | 71.7 ± 3.3 | 7.10 ± 0.07 |
| β-arr1 | 100% | 6.45 ± 0.04 | N/A | N/A | N/A | N/A | N/A | N/A | 16.1 ± 0.7 | 5.95 ± 0.05 |
| β-arr2 | 100% | 6.74 ± 0.07 | 6.3 ± 0.7 | 5.37 ± 0.11 | 24.1 ± 3.8 | 4.92 ± 0.09 | 4.7 ± 0.3 | 5.38 ± 0.16 | 22.4 ± 0.5 | 6.25 ± 0.04 |

| Lofentanil | WT | | Y326F | | I322A | | Q124A | | W293F | |
|---|---|---|---|---|---|---|---|---|---|---|
| | Emax% | pEC50 | Emax% | pEC50 | Emax% | pEC50 | Emax% | pEC50 | Emax% | pEC50 |
| Gi1 | 100% | 9.86 ± 0.06 | 86.8 ± 3.6 | 9.25 ± 0.04 | 101.0 ± 3.7 | 10.04 ± 0.08 | 104.0 ± 3.5 | 9.81 ± 0.08 | 58.3 ± 1.2 | 9.38 ± 0.09 |
| β-arr1 | 100% | 9.25 ± 0.08 | 40.2 ± 1.6 | 8.27 ± 0.08 | 81.4 ± 8.3 | 9.30 ± 0.06 | 68.9 ± 4.9 | 9.13 ± 0.05 | 4.9 ± 0.2 | 9.36 ± 0.07 |
| β-arr2 | 100% | 9.42 ± 0.07 | 43.4 ± 1.2 | 8.55 ± 0.12 | 90.7 ± 3.3 | 9.54 ± 0.09 | 68.6 ± 2.0 | 9.41 ± 0.15 | 7.6 ± 0.2 | 9.59 ± 0.12 |

| MP | WT | | Y326F | | I322A | | Q124A | | W293F | |
|---|---|---|---|---|---|---|---|---|---|---|
| | Emax% | pEC50 | Emax% | pEC50 | Emax% | pEC50 | Emax% | pEC50 | Emax% | pEC50 |
| Gi1 | 100% | 8.34 ± 0.08 | 32.4 ± 0.7 | 6.12 ± 0.02 | 68.2 ± 5.3 | 6.92 ± 0.07 | 39.6 ± 6.4 | 7.32 ± 0.16 | 73.6 ± 2.6 | 7.90 ± 0.04 |
| β-arr1 | 100% | 7.78 ± 0.06 | N/A | N/A | N/A | N/A | N/A | N/A | 148 ± 16 | 7.05 ± 0.04 |
| β-arr2 | 100% | 8.20 ± 0.07 | N/A | N/A | 73 ± 16 | 6.60 ± 0.15 | N/A | N/A | 121 ± 14 | 7.32 ± 0.04 |

**Extended Data Fig.7 | Signaling characterization μOR mutants upon various ligands**

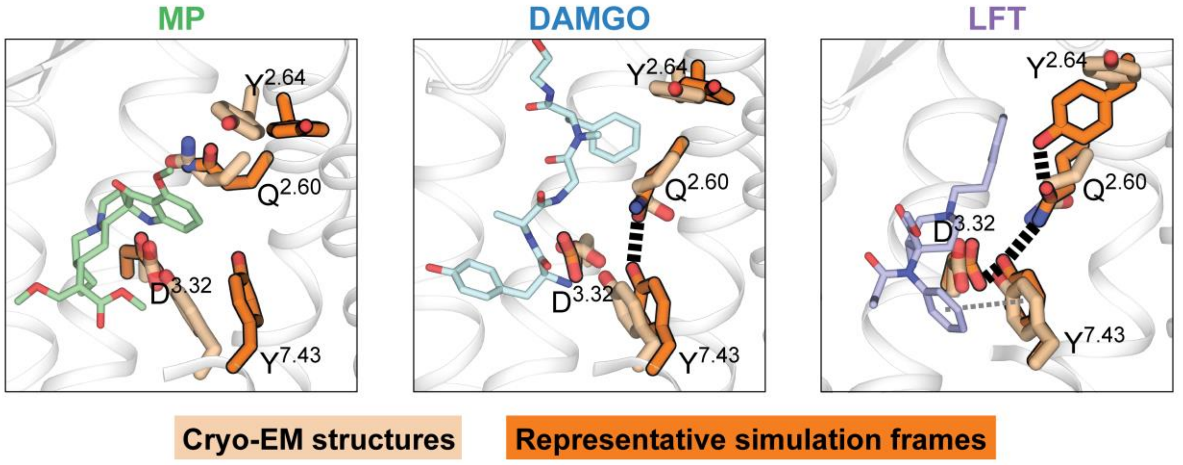


**Extended Data Fig. 8 | Comparison of binding pocket conformations observed in cryo-EM structures and in simulation.**

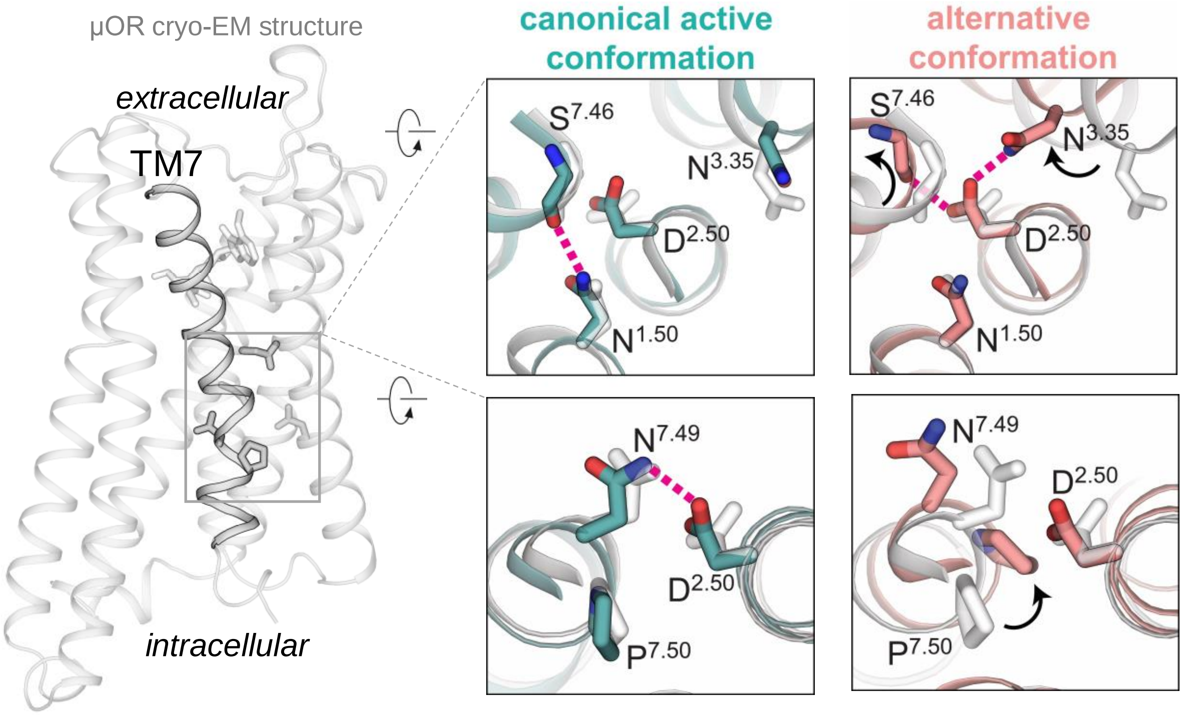


**Extended Data Fig. 9 | The transition from the canonical active conformation to the alternative conformation involves changes in an interaction network in the core of the receptor**.

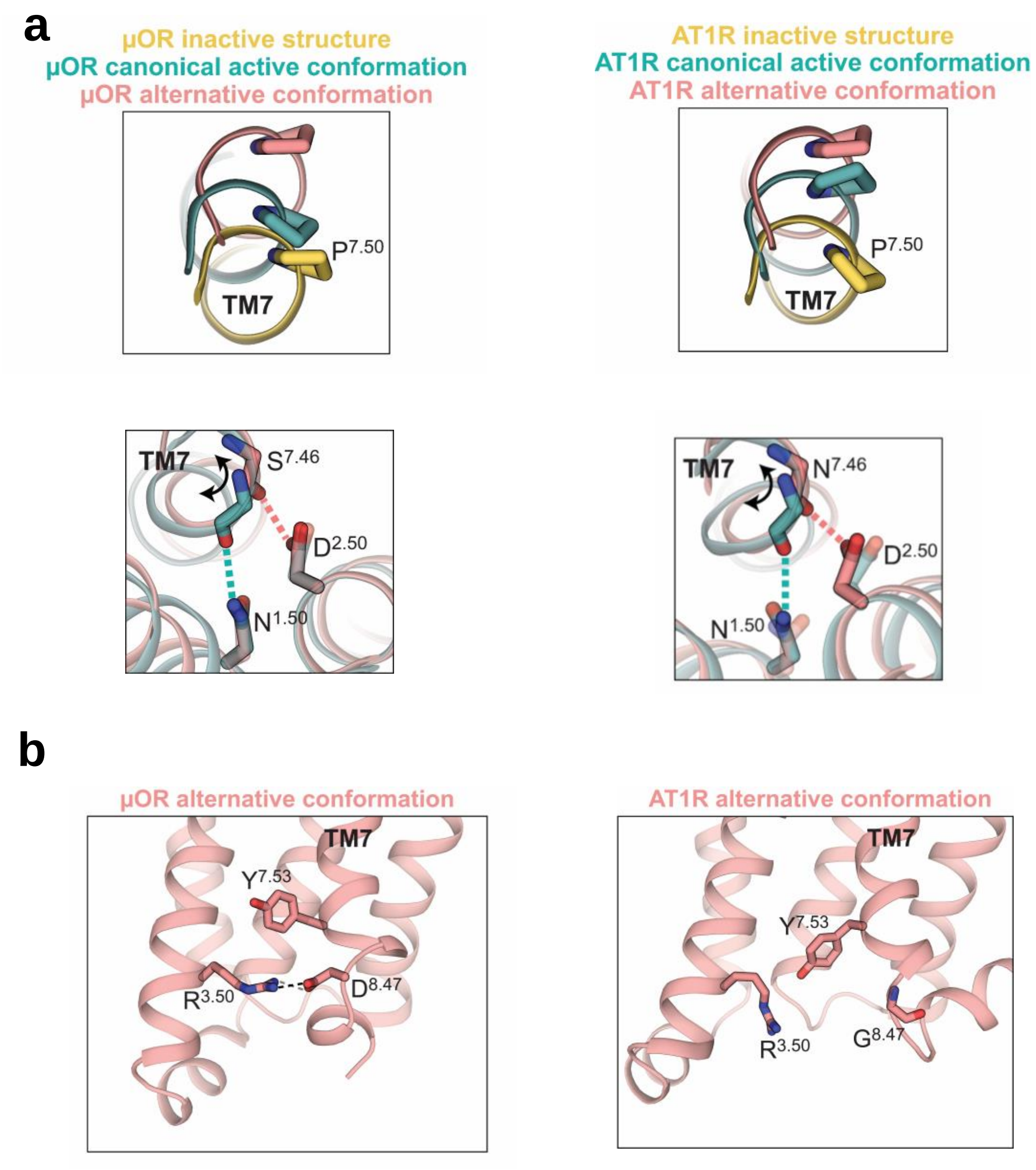


**Extended Data Fig. 10 | Comparison of the intracellular conformations observed for the μOR and the AT1R.**

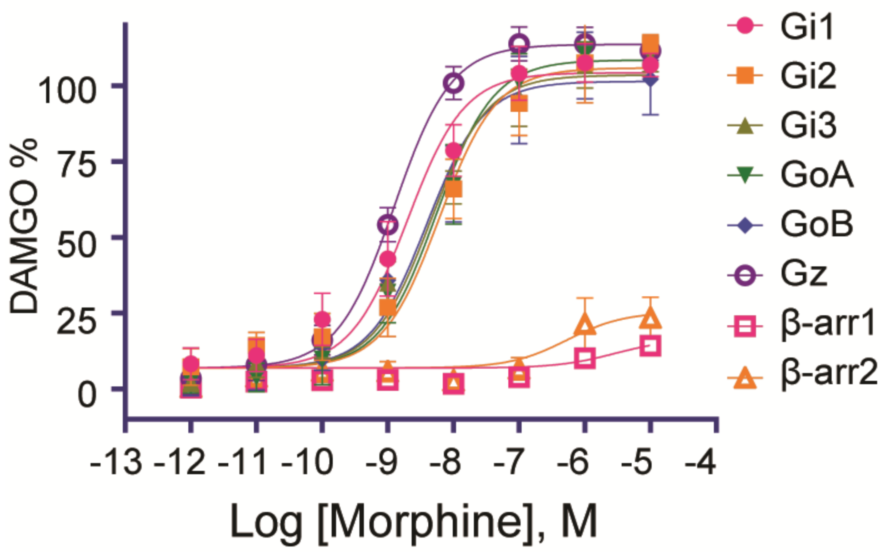


**Extended Data Fig. 11 | Signaling profile of μOR modulated by morphine.**

| MP | | pEC50 | EC50 |
|---|---|---|---|
| | Gi1 | 8.28± 0.14 | 5.30458E-09 |
| | Gi2 | 8.47± 0.12 | 3.40491E-09 |
| | Gi3 | 8.49± 0.15 | 3.24239E-09 |
| | GoA | 8.54± 0.12 | 2.86553E-09 |
| | GoB | 8.27± 0.12 | 5.3769E-09 |
| | Gz | 8.24± 0.21 | 5.82015E-09 |
| | βArrestin1 | N/A | N/A |
| | βArrestin2 | N/A | N/A |
| Lofentanil | | | |
| | Gi1 | 10.04± 0.12 | 9.06418E-11 |
| | Gi2 | 10.14± 0.13 | 7.29902E-11 |
| | Gi3 | 10.04± 0.14 | 9.07695E-11 |
| | GoA | 10.29± 0.12 | 5.16091E-11 |
| | GoB | 10.02± 0.13 | 9.59933E-11 |
| | Gz | 10.22± 0.13 | 6.09333E-11 |
| | βArrestin1 | 10.14± 0.12 | 7.31138E-11 |
| | βArrestin2 | 10.50± 0.15 | 3.18932E-11 |
| DAMGO | | | |
| | Gi1 | 8.08± 0.14 | 8.39532E-09 |
| | Gi2 | 8.12± 0.13 | 7.56742E-09 |
| | Gi3 | 8.11+0.13 | 7.74586E-09 |
| | GoA | 8.01± 0.13 | 9.66749E-09 |
| | GoB | 8.13± 0.13 | 7.40083E-09 |
| | Gz | 8.18± 0.14 | 6.61595E-09 |
| | βArrestin1 | 8.40± 0.14 | 3.9467E-09 |
| | βArrestin2 | 8.71± 0.14 | 1.93612E-09 |

**Extended Data Table 1. Summary of BRET assay results**

| Data collection and processing | | |
|---|---|---|
| | **MP-µOR-Gi** | **LFT-µOR-Gi-scFv** |
| Magnification | 165,000 | 130,000 |
| Voltage (kV) | 300 | 300 |
| Electron exposure (e-/Å$^2$) | 56 | 56 |
| Defocus range (µm) | -0.8 ~-1.8 | -1.0~-2.0 |
| Pixel size (Å) | 0.82 | 1.06 |
| Symmetry imposed | C1 | C1 |
| Final particle images (no.) | 413,821 | 268,658 |
| Map resolution(Å) (Å) | 2.5 | 3.2 |
| FSC threshold | 0.143 | 0.143 |
| **Refinement** | | |
| Initial model used | 6DDF | 6DDE |
| Map sharpening B-factor (Å$^2$) | -66.3 | -126.8 |
| Non-hydrogen atoms | 6860 | 8677 |
| Protein residues | 898 | 1119 |
| Water | 5 | 0 |
| Ligands | 2 | 1 |
| B-factors (Å$^2$) | | |
| Protein | 23.47 | 51.59 |
| Ligand | 14.47 | 62.55 |
| R.m.s. deviations | | |
| Bond lengths (Å) | 0.002 | 0.003 |
| Bond angles (˚) | 0.458 | 0.502 |
| **Validation** | | |
| MolProbity | 1.49 | 1.71 |
| Clashscore | 9.22 | 8.68 |
| Poor rotamers (%) | 0.43 | 0.65 |
| Ramachandran plot | | |
| Favored (%) | 98.09 | 96.37 |
| Allowed (%) | 1.91 | 3.63 |
| Disallowed (%) | 0.00 | 0.00 |

**Extended Data Table 2. Cryo-EM data collection, refinement, and validation statistics**
